# MAP4K3 inhibits Sirtuin-1 to repress the LKB1-AMPK pathway to promote amino acid dependent activation of the mTORC1 complex

**DOI:** 10.1101/2021.11.08.467702

**Authors:** Mary Rose Branch, Cynthia L. Hsu, Kohta Ohnishi, Elian Lee, Jill Meisenhelder, Brett Winborn, Bryce L. Sopher, J. Paul Taylor, Tony Hunter, Albert R. La Spada

## Abstract

mTORC1 is the key rheostat controlling cellular metabolic state. Of the various inputs to mTORC1, the most potent effector of intracellular nutrient status is amino acid supply. Despite an established role for MAP4K3 in promoting mTORC1 activation in the presence of amino acids, the signaling pathway by which MAP4K3 controls mTORC1 activation remains unknown. Here we examined the process of MAP4K3 activation of mTORC1 and found that MAP4K3 represses the LKB1-AMPK pathway to prevent TSC1/2 complex inactivation of Rheb. When we sought the regulatory link between MAP4K3 and LKB1 inhibition, we discovered that MAP4K3 physically interacts with the master nutrient regulatory factor Sirtuin-1 and phosphorylates Sirtuin-1 to repress LKB1 activation. Our results reveal the existence of a novel signaling pathway linking amino acid satiety with MAP4K3-dependent suppression of SIRT1 to inactivate the repressive LKB1-AMPK pathway and thereby potently activate the mTORC1 complex to dictate the metabolic disposition of the cell.

## Introduction

The mechanistic target of rapamycin (mTOR) is a serine/threonine protein kinase that serves as the catalytic subunit of two important regulatory protein complexes, mTORC1 and mTORC2 (Saxton & Sabatini, 2017). While many factors participate in setting the metabolic state of the cell, mTORC1 has emerged as the key rheostat whose activation status determines whether the cell will adopt an anabolic or catabolic state. Both extracellular nutrient status and intracellular nutrient levels provide separate input to mTORC1 via distinct signaling cascades. In times of systemic nutrient abundance, growth factors are released and bind to extracellular receptors which activate the Akt/PI3 kinase pathway to inactivate the TSC1/2 complex, thereby permitting Rheb to remain GTP-bound and active (Wolfson & Sabatini, 2017). As Rheb resides at the surface of the lysosome, which is now known to be the major site of integration of nutrient sensing-dependent cellular signaling, localization of the mTORC1 complex to the lysosome must also occur for full mTORC1 activation. When intracellular nutrients are replete, activation of a set of Rag protein dimers recruits mTORC1 to the lysosome via their interaction with the Ragulator complex (Lehman & Abraham, 2020). Activation of the mTORC1 complex promotes protein synthesis as well as de novo lipid and nucleotide synthesis, and suppresses catabolic processes, including especially autophagy (Saxton & Sabatini, 2017).

Although multiple inputs coalesce on the mTORC1 complex via various, different signaling pathways, the most potent determinant of intracellular nutrient status is amino acid supply, reflecting nitrogen levels (Smith *et al*, 2005; Yan & Lamb, 2012). The cell is especially attuned to the levels of certain essential amino acids, such as leucine. Regulation of the Rag proteins, which recruit mTORC1 to the lysosome, is dictated by the GATOR1 and GATOR2 complexes, which are responsive to the cellular amino acid sensor Sestrin-2 and its related family members (Wolfson & Sabatini, 2017). When leucine is abundant, Sestrin-2 is bound by leucine, and this leucylation of Sestrin-2 causes dissociation from GATOR2, thereby relieving GATOR2 inhibition; once active, GATOR2 represses GATOR1 inhibition of the Rag proteins, resulting in Rag-dependent recruitment of mTORC1 to the surface of the lysosome.

Mitogen-activated protein kinases (MAPKs) comprise a large family of key regulatory proteins that control a broad range of essential processes in eukaryotic cells (Qi & Elion, 2005). MAP4K3, also known as germinal-center kinase-like kinase (GLK), is a member of the Ste20 sub-family of MAPKs (Diener *et al*, 1997). Studies in mammalian cell lines and in *Drosophila* have shown that MAP4K3 is absolutely required for activation of mTORC1 in response to amino acids (Bryk *et al*, 2010; Findlay *et al*, 2007; Resnik-Docampo & de Celis, 2011). Furthermore, MAP4K3 is ubiquitously expressed, as MAP4K3 RNA and protein are detected in all human tissues (Diener *et al*., 1997; Uhlen *et al*, 2015). Thus, MAP4K3 likely has a central role in regulating the metabolic disposition of the cell.

Nitrogen status is central to the cell’s decision to adopt a catabolic or anabolic state. While studying the transcriptional regulation of autophagy, we discovered that microRNA *let-7* can activate autophagy by coordinately down-regulating the expression of genes whose protein products mediate amino acid dependent activation of the mTORC1 complex, a potent repressor of autophagy (Dubinsky *et al*, 2014). We identified MAP4K3 as one such target, and we documented that knock-down of MAP4K3 alone is sufficient to robustly induce autophagy (Dubinsky *et al*., 2014). To determine the mechanistic basis for MAP4K3 regulation of autophagy, we considered key nodes involved in dictating the status of the autophagy pathway in the cell. Transcription factor EB (TFEB) is a helix-loop-helix transcription factor that localizes to the nucleus under conditions of nutrient depletion or cellular stress to drive the expression of a suite of genes necessary for autophagosome formation, autophagosome-lysosome fusion, and lysosome formation and function (Sardiello *et al*, 2009). Although it was known that mTORC1 represses TFEB and restricts it to the cytosol by phosphorylating it at serine 211 (Martina *et al*, 2012; Roczniak-Ferguson *et al*, 2012; Settembre *et al*, 2012), we found that this regulatory activity is dependent upon MAP4K3. Specifically, we found that MAP4K3 must first phosphorylate TFEB at serine 3 in order for mTORC1 to phosphorylate TFEB at serine 211 (Hsu *et al*, 2018). Our results established that MAP4K3 lies upstream of mTORC1 in the negative regulation of autophagy, suggesting that MAP4K3 is likely a central nutrient-sensing regulator in the cell.

Despite a role for MAP4K3 in promoting cell proliferation in the presence of abundant amino acids, the signaling pathway by which MAP4K3 controls mTORC1 activation remains unknown. Here we examined the process of MAP4K3 activation of mTORC1 when the cell is stimulated with amino acids, and we found that impaired mTORC1 activation upon loss of MAP4K3 can be rescued by elimination of AMPK or LKB1, suggesting that MAP4K3 activation of mTORC1 operates via the LKB1-AMPK pathway to prevent TSC1/2 complex inactivation of Rheb. When we sought the regulatory link between MAP4K3 and inhibition of LKB1, we discovered that MAP4K3 directly interacts with the master nutrient regulatory deacetylase Sirtuin-1 (SIRT1) and phosphorylates SIRT1 on threonine 344 (T344) to prevent LKB1 deacetylation and thereby repress LKB1 activation. Our results reveal the existence of a novel signaling pathway linking amino acid satiety with MAP4K3-dependent suppression of SIRT1 to inactivate the repressive LKB1-AMPK pathway and thereby potently activate the mTORC1 complex to dictate the metabolic disposition of the cell.

## Results

### MAP4K3 activation of mTORC1 in the presence of amino acids is Rheb-dependent

To determine the role of MAP4K3 in regulating cell growth, we compared the proliferation of wild-type HEK293A cells with two independently derived clonal MAP4K3 HEK293A knock-out (k.o.) cell lines, which we had previously generated using CRISPR-Cas9 genome editing (Hsu *et al*., 2018). We documented markedly reduced cell growth under nutrient replete conditions in both MAP4K3 k.o. cell lines (**Figure 1A**), and noted that MAP4K3 k.o. cells are also smaller in size (**Figure S1A**). Studies in mammalian cell lines and in *Drosophila* have shown that MAP4K3 is essential for activation of mTORC1 in response to amino acids (Bryk *et al*., 2010; Findlay *et al*., 2007; Resnik-Docampo & de Celis, 2011). Activation of mTORC1 increases protein translation through activation of S6 kinase 1 (S6K) and inhibition of 4E binding protein 1 (4E-BP1) by phosphorylation of these targets, and activated S6K in turn phosphorylates the S6 protein to turn it on (Lipton & Sahin, 2014). When we evaluated the phosphorylation status of S6K, S6, and 4E-BP1 in wild-type (WT) HEK293A cells and in MAP4K3 k.o. cells, we confirmed that MAP4K3 is required for full mTORC1 activation upon amino acid stimulation (**Figures 1B & S1B-C**). Both extracellular nutrient status and intracellular nutrient levels provide separate input to mTORC1 via distinct signaling cascades (Wolfson & Sabatini, 2017). To delineate the pathway by which MAP4K3 activates mTORC1, we evaluated Rheb activity in MAP4K3 k.o. cells, and found that overexpression of the constitutively active Rheb Q641 mutant can activate mTORC1 in MAP4K3 k.o. cells, regardless of amino acid status (**Figure 1C**). This finding indicates that MAP4K3 normally activates mTORC1 through Rheb activation, as Rheb is regulated by the tuberous sclerosis 2 protein (TSC2), which acts as a GTPase-activating protein (GAP) for Rheb (Garami *et al*, 2003; Inoki *et al*, 2003; Tee *et al*, 2003; Zhang *et al*, 2003). As TSC2 preferentially interacts with GTP-bound Rheb (Carroll *et al*, 2016), we documented that interaction between endogenous TSC2 and Rheb in WT cells was much stronger than in MAP4K3 k.o. cells under conditions of nutrient satiety (**Figure 1D**). This is indicative of a markedly higher proportion of inactive GDP-bound Rheb when MAP4K3 is turned off or absent. When we considered the amino acid sensing requirements for MAP4K3 activation, we found that refeeding starved WT and MAP4K3 k.o. cells with either leucine or arginine yielded reduced mTORC1 activation in both cases (**Figure S1D-E**), suggesting that MAP4K3 activation is not restricted to a particular amino acid.

**Figure 1.**
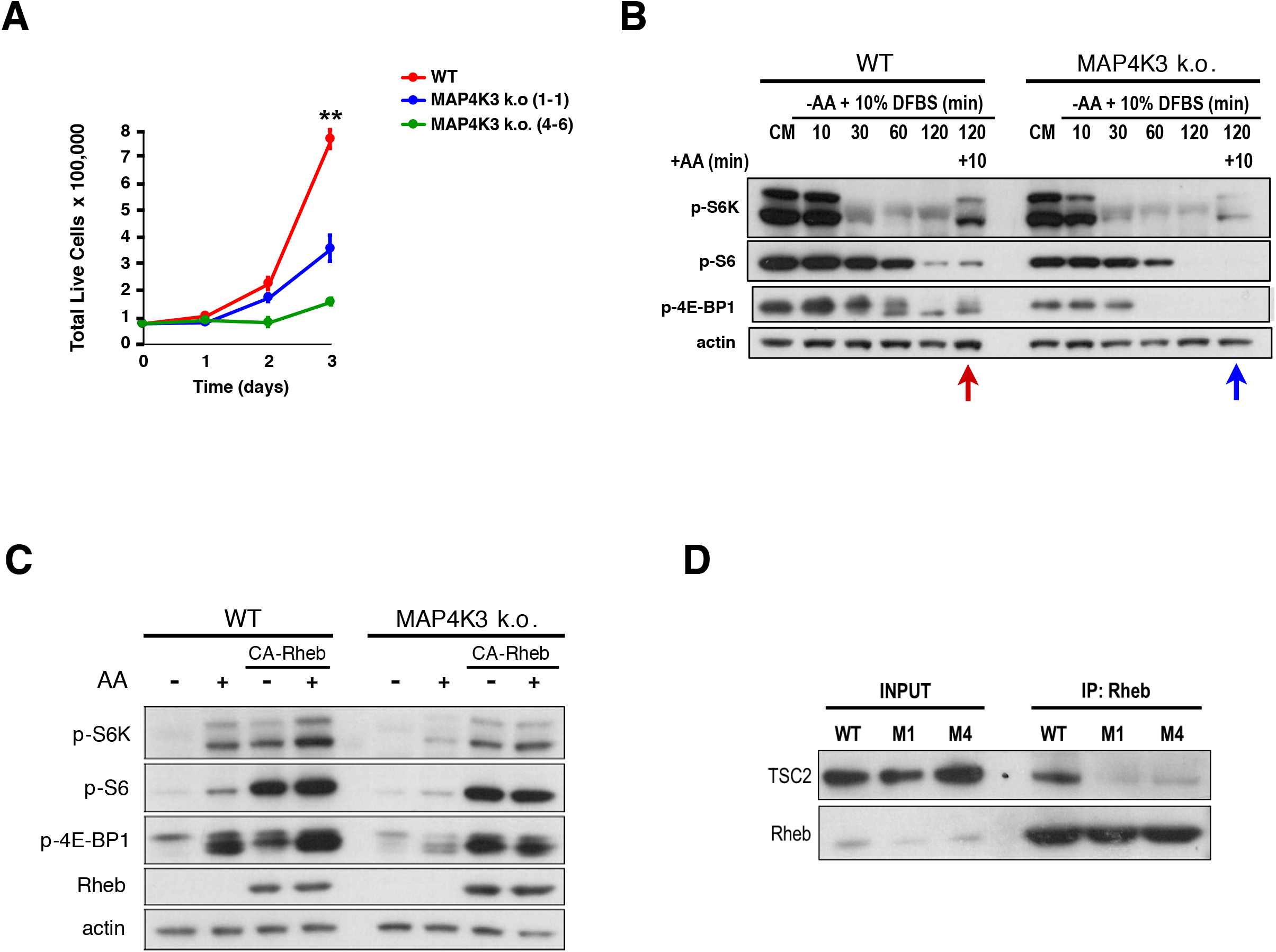
MAP4K3 is required for Rheb-dependent activation of mTORC1 in the presence of amino acids. **(A)** We cultured a WT HEK293A cell line and two uniquely generated MAP4K3 knock-out (k.o.) cell lines in complete media over 72 hrs. Here we see quantification of cell numbers at 24 hr intervals. \*\**P* < 0.01, ANOVA with post-hoc Tukey test; n = 3 biological replicates. Error bars = s.e.m. **(B)** WT and MAP4K3 k.o. cells were starved for 10, 30, 60, and 120 min, and then re-stimulated with amino acids for 10 min. We immunoblotted resulting cell protein lysates for phosphorylated S6 kinase 1, phosphorylated S6, and phosphorylated 4E-BP1, as indicated. Note reduced phosphoactivation of these mTORC1 targets in the MAP4K3 k.o. cells (blue arrow) compared to WT cells (red arrow). β-actin served as the loading control. See Figure S1 for measurement of mTORC1 phospho-targets relative to corresponding total protein with quantification. **(C)** WT and MAP4K3 k.o. cells were transfected with constitutively active Rheb where indicated, and starved of amino acids for 3 hrs (-), or starved of amino acids for 3 hrs and then re-stimulated with amino acids for 10 min (+). Protein lysates were prepared and immunoblotted for the indicated proteins. β-actin served as the loading control. **(D)** Protein lysates were prepared from a WT HEK293A cell line and two uniquely generated MAP4K3 k.o. cell lines, and immunoprecipitated with anti-Rheb antibody for endogenous TSC2. Note minimal interaction of Rheb with TSC2 in both MAP4K3 k.o. cell lines.

### MAP4K3 represses the LKB1 - AMPK pathway to activate the mTORC1 complex

There are two major inputs to the TSC1/2 complex upstream of Rheb: i) the phosphatidyl-inositol 3-kinase - Akt / protein kinase B (PI3K-Akt) pathway; and ii) adenosine monophosphate activated protein kinase (AMPK). To determine if MAP4K3 activation of mTORC1 operates through either of these pathways, we examined the activation status of Akt and AMPK in MAP4K3 k.o. cells, and noted that while Akt phospho-activation was not different between WT and MAP4K3 k.o. cells (**Figure 2A**), AMPKα1 subunit phospho-activation was markedly increased in MAP4K3 k.o. cells (**Figure S2A**), and AMPK inhibitory phosphorylation of acetyl-CoA carboxylase (ACC), one of its main targets, was also correspondingly increased in MAP4K3 k.o. cells upon amino acid stimulation (**Figure 2B**). To test the hypothesis that MAP4K3 is acting upstream of AMPK to activate mTORC1, we used CRISPR-Cas9 genome editing to generate MAP4K3/AMPKα1 double k.o. cells. When we compared cell proliferation between WT, MAP4K3 k.o., AMPKα1 k.o., and MAP4K3/AMPKα1 double k.o. cells, we found that the reduced cell growth phenotype observed in MAP4K3 k.o. cells was markedly improved in MAP4K3/AMPKα1 double k.o. cells (**Figures 2C & S2B**), and noted that MAP4K3/AMPKα1 double k.o. cells are of normal size (**Figure S2C**). Furthermore, upon amino acid stimulation, MAP4K3/AMPKα1 double k.o. cells exhibited robust mTORC1 activation (**Figure 2D**), indicating that concomitant absence of AMPK could fully rescue the mTORC1 inhibition occurring in MAP4K3 k.o. cells. Although MAP4K3 is required to achieve full mTORC1 activation upon amino acid stimulation, MAP4K3 is not necessary for mTORC1 activation when MAP4K3 k.o. cells are grown in the presence of glucose (**Figure S2D**). AMPK, however, promotes mTORC1 repression upon glucose starvation, and concomitant absence of MAP4K3 does not affect mTORC1 activation status under these conditions (**Figure S2D**).

**Figure 2.**
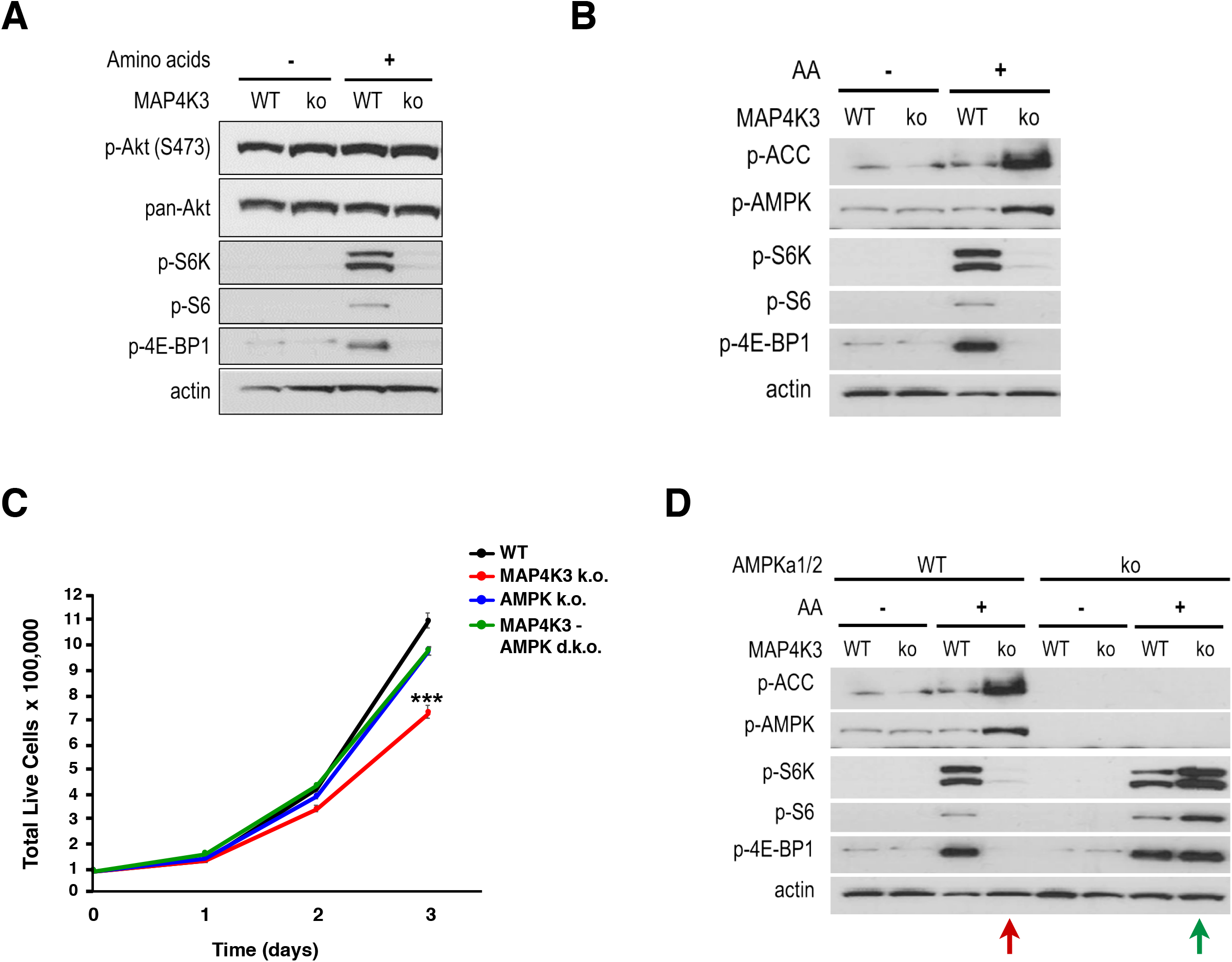
Activated MAP4K3 represses AMPK to turn on the mTORC1 complex. **(A)** WT and MAP4K3 k.o. cells were starved of amino acids for 3 hrs (-), or starved of amino acids for 3 hrs and then re-stimulated with amino acids for 10 min (+). We immunoblotted resulting cell protein lysates for phosphorylated Akt, total Akt, phosphorylated S6 kinase 1, phosphorylated S6, and phosphorylated 4E-BP1, as indicated. β-actin served as the loading control. **(B)** WT and MAP4K3 k.o. cells were starved of amino acids for 3 hrs (-), or starved of amino acids for 3 hrs and then re-stimulated with amino acids for 10 min (+). We immunoblotted resulting cell protein lysates for phosphorylated ACC, phosphorylated AMPK α1 subunit, phosphorylated S6 kinase 1, phosphorylated S6, and phosphorylated 4E-BP1, as indicated. β-actin served as the loading control. See Figure S2 for measurement of phosphorylated AMPK α1 subunit relative to corresponding total protein. **(C)** We cultured a WT HEK293A cell line, MAP4K3 k.o. cell line, AMPK α1 k.o. cell line, and MAPK3/AMPK α1 double k.o. cell line in complete media over 72 hrs. Here we see quantification of cell numbers at 24 hr intervals. \*\*\**P* < 0.001; ANOVA with post-hoc Tukey test. n = 3 technical replicates. Error bars = s.e.m. See Figure S2 for bar graph of terminal cell number data. **(D)** WT cells, MAP4K3 k.o. cells, AMPK α1 k.o. cells, and MAPK3/AMPK α1 double k.o. cells were starved of amino acids for 3 hrs (-), or starved of amino acids for 3 hrs and then re-stimulated with amino acids for 10 min (+). We immunoblotted resulting cell protein lysates for phosphorylated ACC, phosphorylated AMPK, phosphorylated S6 kinase 1, phosphorylated S6, and phosphorylated 4E-BP1, as indicated. Note complete rescue of mTORC1 activation in the MAP4K3/AMPK α1 double k.o. cell line. β-actin served as the loading control.

A series of studies in worms, flies, and mice have established that Liver Kinase B1 (LKB1) is the major upstream regulator of AMPK, and that LKB1 phosphoactivation of AMPK leads to mTORC1 repression (Kullmann & Krahn, 2018). To determine if MAP4K3 activation of mTORC1 is LKB1 dependent, we examined the subcellular localization of LKB1, as translocation of LKB1 out of the nucleus into the cytosol is required for LKB1 activation of AMPK. When we compared LKB1 translocation to the cytosol in WT HEK293A cells and MAP4K3 k.o. HEK293A cells, we observed a marked reduction of LKB1 in the cytosol of WT cells subjected to amino acid refeeding; however, such amino acid stimulation did not prevent LKB1 translocation into the cytosol in cells lacking MAP4K3 (**Figure 3A-B**). Another indicator of LKB1 activation status is Sirtuin-1 dependent deacetylation (Lan *et al*, 2008). Using the same paradigm of amino acid depletion and refeeding, we measured a >50% reduction in LKB1 acetylation upon amino acid starvation in WT cells, which is consistent with LKB1 activation, and then detected a nearly 3-fold increase in LKB1 acetylation levels in WT cells upon amino acid refeeding, indicative of LKB1 inhibition upon nutrient satiety (**Figure 3C**). In MAP4K3 k.o. cells, LKB1 acetylation levels increased less than 2-fold upon amino acid stimulation, suggesting only partial inhibition in the absence of MAP4K3 (**Figure 3C**). To clarify the role of LKB1 in MAP4K3 amino-acid dependent activation of mTORC1, we used CRISPR-Cas9 genome editing to derive MAP4K3/LKB1 double k.o. cells. Similar to the results obtained with the MAP4K3/AMPK double k.o. cells, we found that MAP4K3/LKB1 double k.o. cells displayed robust mTORC1 activation in response to amino acid stimulation, while MAP4K3 single k.o. cells exhibited no detectable evidence of mTORC1 activation in the presence of amino acids (**Figure 3D**). These results indicate that MAP4K3 is acting upstream of the LKB1-AMPK axis to activate the mTORC1 complex.

**Figure 3.**
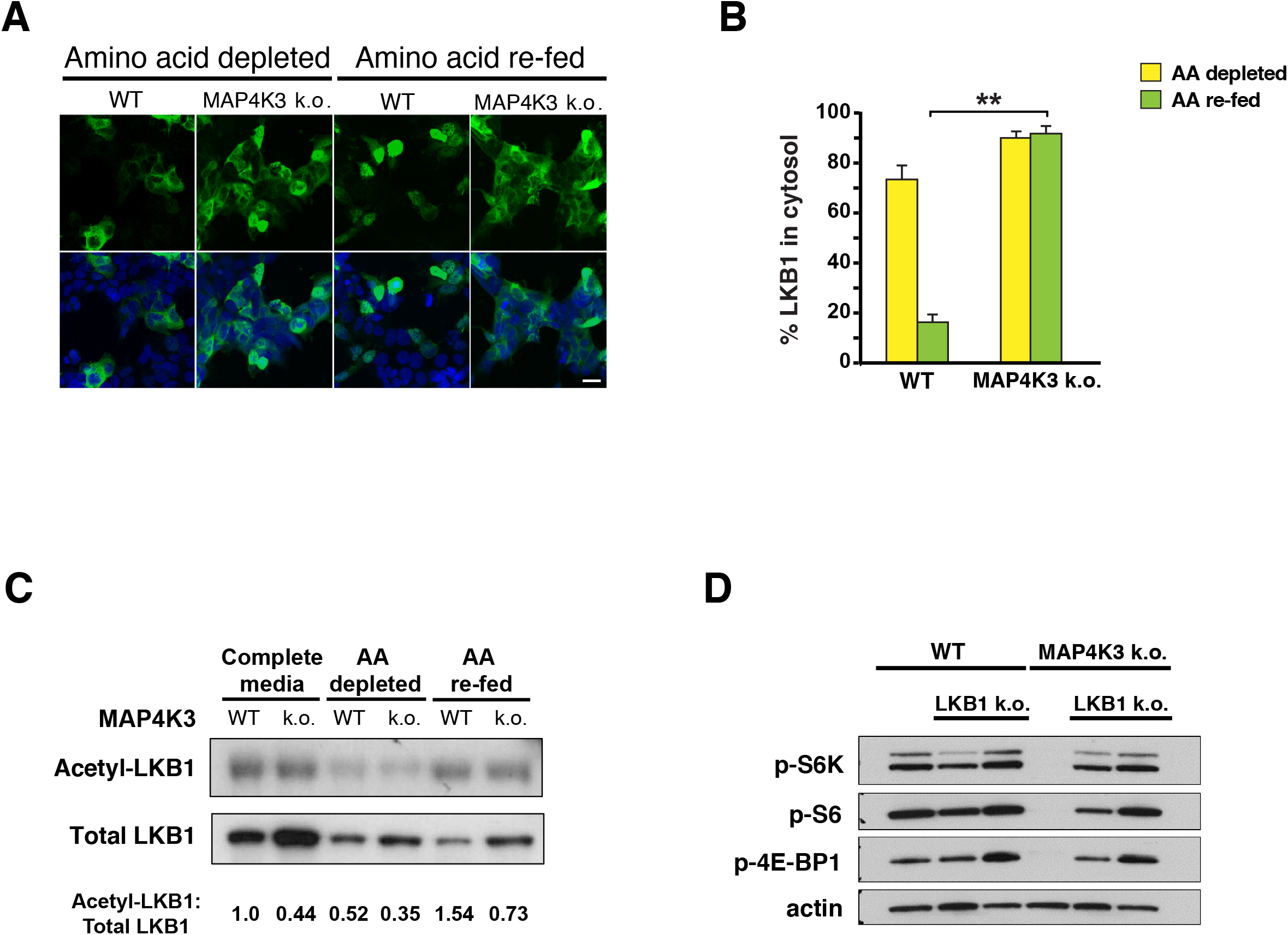
MAP4K3 amino acid dependent activation of mTORC1 occurs via LKB1 repression. **(A)** WT and MAP4K3 k.o. cells were transfected with a LKB1-FLAG vector, and then starved of amino acids for 3 hrs (= amino acid depleted), or starved of amino acids for 3 hrs and re-stimulated with amino acids for 10 min (= amino acid re-fed). Cells were fixed, immunostained with anti-FLAG antibody (green) and counterstained with DAPI (blue). Scale bar, 10 μm. **(B)** Quantification of the percentage of cells showing LKB1 localization to the cytosol in **(A)**. Note that LKB1 is not retained in the nucleus in MAP4K3 k.o. cells upon amino acid stimulation. *P* < 0.01, two-tailed t-test; n = 3 biological replicates. Error bars = s.e.m. **(C)** WT and MAP4K3 k.o. cells were transfected with a LKB1-FLAG vector, and then starved of amino acids for 3 hrs (= AA depleted), or starved of amino acids for 3 hrs and re-stimulated with amino acids for 10 min (= AA re-fed). We prepared protein lysates and performed anti-FLAG immunoprecipitation (IP) of LKB1, and immunoblotted IP material with an anti-acetyl lysine antibody or an anti-LKB1 antibody. After densitometry analysis, we calculated the ratio of acetylated LKB1: total LKB1. Results shown are normalized to WT cells in complete media, which was arbitrarily set to 1. **(D)** WT cells, MAP4K3 k.o. cells, LKB1 k.o. cells, and MAPK3/LKB1 double k.o. cells were starved of amino acids for 3 hrs and then re-stimulated with amino acids for 30 min. We immunoblotted resulting cell protein lysates for phosphorylated S6 kinase 1, phosphorylated S6, and phosphorylated 4E-BP1, as indicated. Note complete rescue of mTORC1 activation in the MAP4K3/LKB1 double k.o. cell line. β-actin served as the loading control.

### MAP4K3 interacts with and phosphorylates Sirtuin-1

To identify MAP4K3 interacting proteins potentially involved in the upstream regulation of mTORC1 activation, we performed mass spectrometry on HeLa cells transiently transfected with FLAG-tagged MAP4K3, and we found that Sirtuin-1 (SIRT1) was among the MAP4K3 interactors (**Data file 1**). As SIRT1 is a NAD^+^-dependent deacetylase that has been reported to activate LKB1 via deacetylation (Lan *et al*., 2008), we hypothesized that MAP4K3 repression of the LKB1-AMPK axis may occur through inhibition of SIRT1. To test this hypothesis, we performed co-transfection co-immunoprecipitation studies in HEK293A cells, and detected a physical interaction between MAP4K3 and SIRT1 (**Figure 4A**). We noted that the interaction between kinase-dead (KD)-MAP4K3 and SIRT1 was stronger than the interaction between WT-MAP4K3 and SIRT1 (**Figure 4A**), indicating that the interaction may depend upon the kinase activity of MAP4K3. To further explore the nature of their physical interaction, we produced SIRT1 by in vitro transcription-translation, and performed pull-down assays with WT-MAP4K3, KD-MAP4K3, or FLAG-GFP empty vector. While we failed to detect a physical interaction between FLAG-tagged GFP and HA-tagged SIRT1, we observed evidence for an interaction between FLAG-tagged MAP4K3 and HA-tagged SIRT1, noting a slightly increased interaction between KD-MAP4K3 and SIRT1 (**Figure 4B**). To directly examine if MAP4K3 is capable of phosphorylating SIRT1, we performed phosphopeptide mapping of SIRT1 isolated from P^32^-labeled WT HEK293A cells and MAP4K3 k.o. cells in the absence or presence of amino acids, and we observed amino acid-dependent phosphopeptide fragments of SIRT1 that were present only in WT cells, and based upon mobility predictions, these fragments were consistent with peptides containing threonine phosphorylation (**Figure 4C**). To help identify likely SIRT1 amino acid residues subject to phosphorylation by MAP4K3, we performed phosphoamino acid analysis of SIRT1 isolated from WT HEK293A cells subjected to amino acid refeeding, and we resolved phosphoamino acids by two-dimensional gel electrophoresis on thin layer cellulose plates, which revealed an increase in the phosphothreonine content of SIRT1 upon amino acid refeeding (**Figure 4D**).

**Figure 4.**
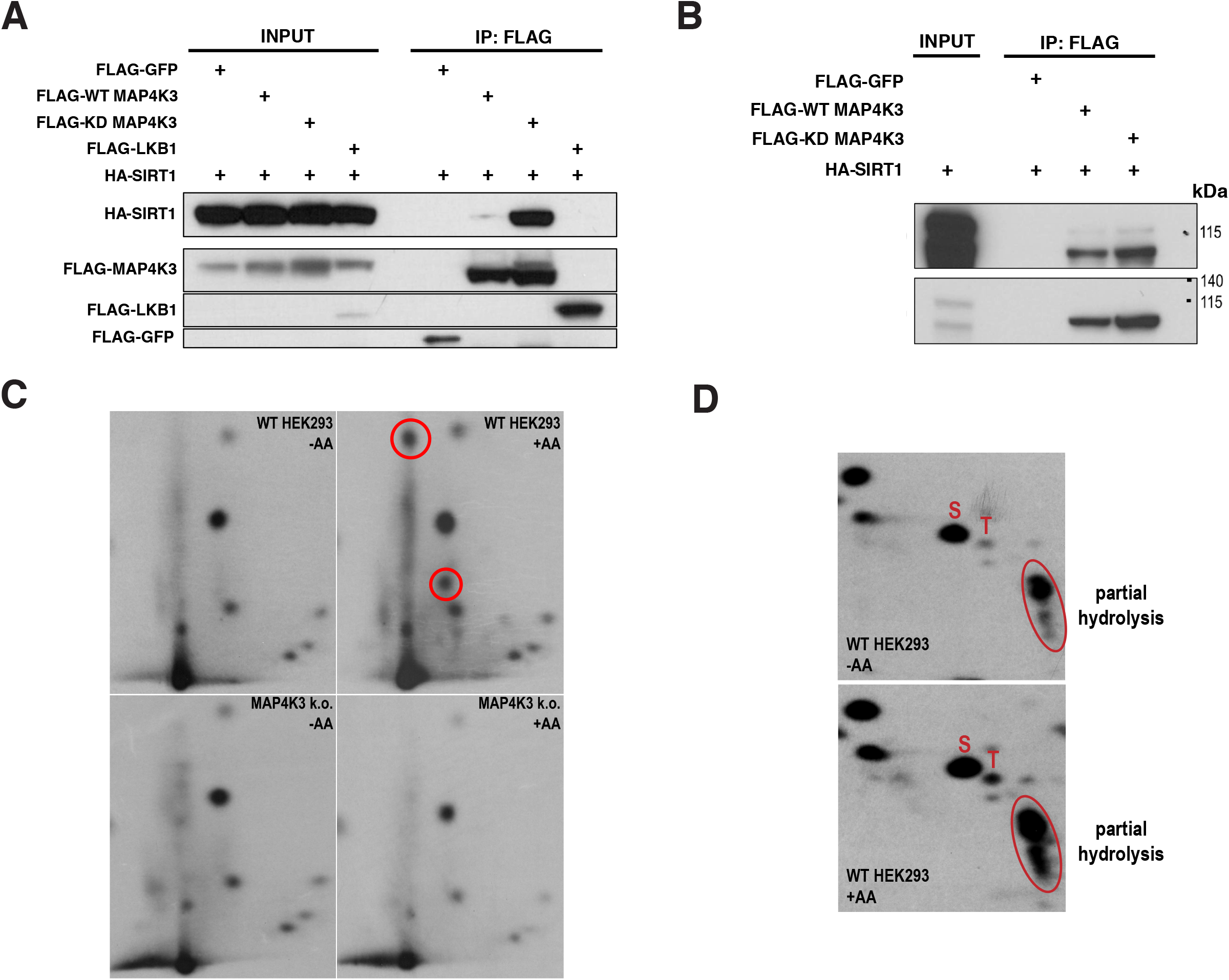
MAP4K3 interacts with and phosphorylates SIRT1. **(A)** HEK293A cells were transfected with SIRT1-HA and WT-MAP4K3-FLAG, kinase dead (KD)-MAP4K3-FLAG, FLAG-GFP empty vector (negative control), or FLAG-LKB1 (positive control). After performing immunoprecipitation (IP) with anti-FLAG antibody, we immunoblotted the IP material for the indicated proteins. **(B)** We incubated in vitro transcribed-translated SIRT1-HA with anti-FLAG IP material isolated from cells expressing either WT-MAP4K3-FLAG, KD-MAP4K3-FLAG, or FLAG-GFP empty vector. We then performed FLAG pull-downs, and immunoblotted with anti-HA antibody and anti-FLAG antibody, which confirmed a physical interaction between recombinant SIRT1 and MAP4K3. **(C)** We transfected WT or MAP4K3 k.o. HEK293A cells with SIRT1-HA, allowed them to grow in labelling media with P^32^ orthophosphate, and subjected the cells to amino acid starvation (-AA) or amino acid starvation followed by amino acid restimulation (+AA). SIRT1 was immunoprecipitated using anti-HA and resolved by SDS-PAGE and autoradiography. SIRT1 protein was isolated, digested with glutamyl endopeptidase, and the resulting peptide mix was spotted on cellulose thin layer plates, for separation by electrophoresis followed by ascending chromatography, and then autoradiography to visualize the phosphopeptides. Circles indicate MAP4K3-dependent threonine phosphorylated peptide fragments of SIRT1. **(D)** After isolating SIRT1 as in ‘C’, SIRT1 was acid hydrolyzed and after mixing with unlabeled phosphoamino acid standards, the amino acid mixtures were separated by two-dimensional electrophoresis on thin layer chromatography plates followed by autoradiography to visualize the P^32^-labeled phosphoamino acids. Unlabeled phosphoamino acid standards were visualized by spraying the thin layer chromatography plates with ninhydrin, and autoradiography films were aligned with the plates to identify the P^32^-labeled phosphoamino acids from the SIRT1 samples. S = phosphoserine, and T = phosphothreonine. Products of partial hydrolysis are circled.

### Sirtuin-1 inhibition is necessary for MAP4K3-dependent activation of mTORC1

If MAP4K3 activation of mTORC1 requires SIRT1 inhibition, then over-expression of SIRT1 in WT cells should blunt the ability of amino acid satiety to turn on the mTORC1 complex. To test this hypothesis, we transfected WT HEK293A cells growing in complete media (CM) with SIRT1-HA or pcDNA empty vector, and then subjected the transfected cells to amino acid starvation, followed by amino acid refeeding. Localization of SIRT1-HA to the nucleus was confirmed by immunostaining (**Figure S3**). As expected, WT HEK293A cells transfected with empty vector displayed repression of mTORC1 activity upon amino acid starvation followed by robust activation of the mTORC1 complex upon amino acid refeeding (**Figure 5A**). However, WT cells over-expressing SIRT1 displayed only about half-maximal mTORC1 complex activation upon amino acid stimulation, when compared to WT cells transfected with empty vector (**Figure 5A-B**). These results indicate that increased levels of SIRT1 may overwhelm MAP4K3’s capacity to repress all of the SIRT1 protein present in the cell, resulting in partial suppression of mTORC1 activation in WT cells upon amino acid stimulation.

**Figure 5.**
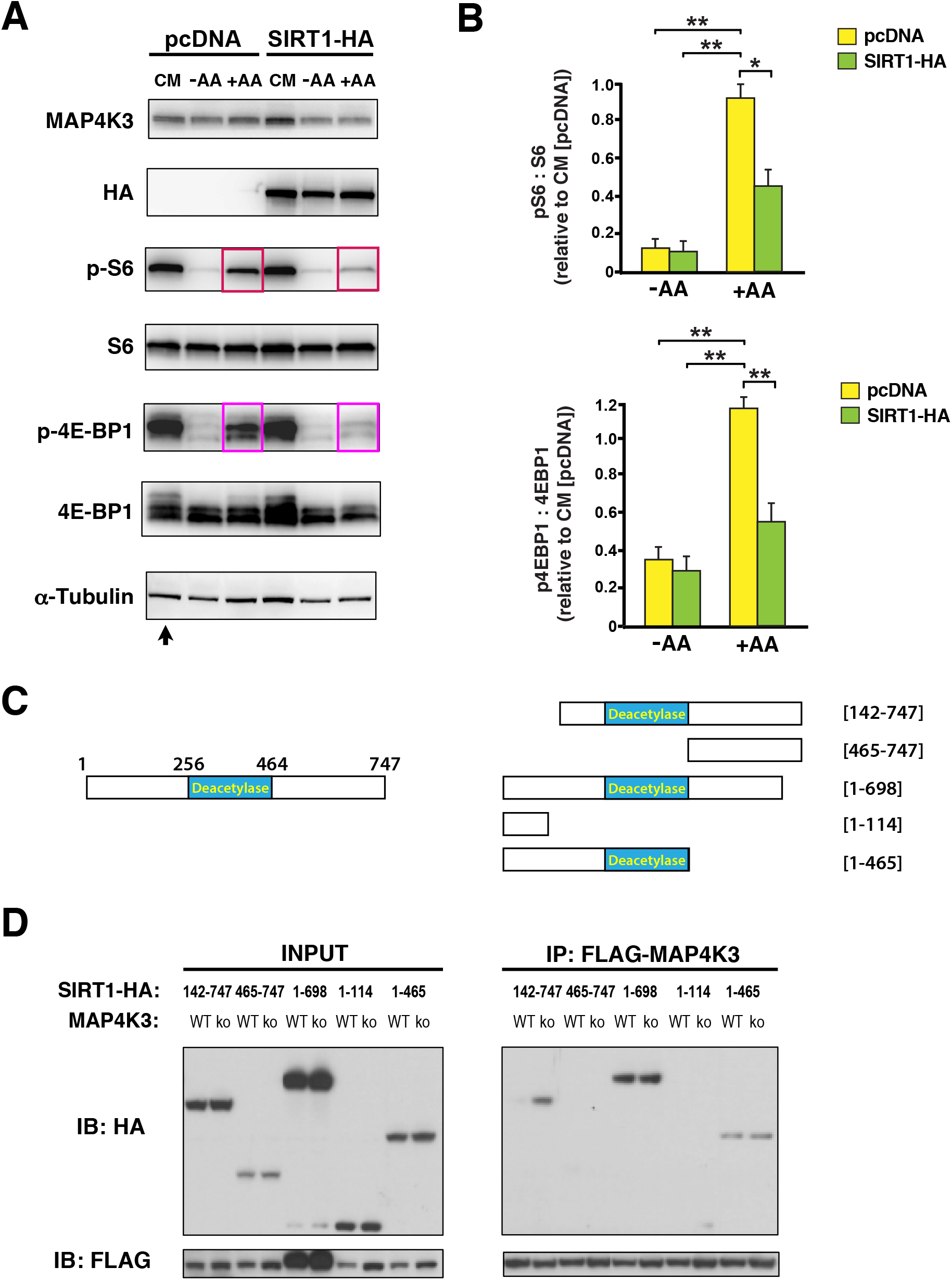
SIRT1 over-expression blunts MAP4K3 amino acid dependent activation of mTORC1 and deletion mapping of SIRT1 interaction with MAP4K3. **(A)** We transfected WT HEK293A cells with either SIRT1-HA or pcDNA empty vector, and maintained the cells under baseline complete media (CM) conditions, or subjected the cells to amino acid starvation (-AA) or amino acid starvation followed by amino acid restimulation (+AA). We immunoblotted resulting cell protein lysates for MAP4K3, HA-tagged SIRT1, phosphorylated S6, total S6, phosphorylated 4E-BP1, and total 4E-BP1, as indicated. Note suppression of mTORC1 activation in WT cells over-expressing SIRT1. α-tubulin served as the loading control. **(B)** TOP GRAPH: We quantified the levels of phosphorylated S6 and total S6 in **(A)** by densitometry, determined the ratio of phosphorylated S6: total S6, and normalized the results to pcDNA-transfected WT HEK293A cells at baseline (arrow in **(A)**). \*\**P* < 0.01, ANOVA with post-hoc Tukey test. BOTTOM GRAPH: We quantified the levels of phosphorylated 4E-BP1 and total 4E-BP1 in **(A)** by densitometry, determined the ratio of phosphorylated 4E-BP1: total 4E-BP1, and normalized the results to pcDNA-transfected WT HEK293A cells at baseline (arrow in **(A)**). \*\**P* < 0.01, ANOVA with post-hoc Tukey test; n = 4 biological replicates. Error bars = s.e.m. **(C)** Diagram of SIRT1 deletion constructs used for mapping its interaction with MAP4K3. LEFT: Full-length SIRT1 with enzymatic deacetylase domain indicated. RIGHT: Diagrams of the different SIRT1 deletion constructs used in the co-transfection, co-immunoprecipitation studies. **(D)** We co-transfected HEK293A cells with either WT-MAP4K3-FLAG or kinase dead (KD)-MAP4K3-FLAG and with a different SIRT1-HA deletion construct, as indicated. We then performed co-immunoprecipitation of MAP4K3 and SIRT1 by FLAG IP, followed by immunoblotting with antiHA antibody or anti-FLAG antibody. Immunoblotting of protein lysates from input cells is shown on the left.

### MAP4K3 phosphorylation of Sirtuin-1 at threonine 344 promotes mTORC1 activation

To identify SIRT1 amino acid regulatory sites subject to MAP4K3 phosphorylation, we pursued two independent lines of investigation. First, we performed phosphoproteomics by transfecting WT HEK293A cells and MAP4K3 k.o. cells with SIRT1-HA, isolated SIRT1-HA protein, and after trypsin digestion, performed mass spectrometry analysis (**Data file 2**). Comparison of phosphopeptides in WT HEK293A and MAP4K3 k.o. cells revealed 14 serine phosphorylation sites and 3 threonine phosphorylation sites that were enriched for SIRT1 isolates from WT cells (**Table S1**). Second, we transfected WT HEK293A cells with FLAG-tagged WT-MAP4K3 or KD-MAP4K3 in combination with different fragments of HA-tagged SIRT1 (**Figure 5C**), and then performed co-immunoprecipitations with anti-FLAG antibody. We detected interactions between MAP4K3 and SIRT1:142-747, SIRT1:1-465, and SIRT1:l-698, but not between MAP4K3 and SIRT1:1-114 or SIRT1:465-747 (**Figure 5D**). When we considered the results of three different independent lines of investigation - phosphoamino acid analysis (**Figure 4C-D**), phosphoproteomics mass spectrometry (**Table S1**), and MAP4K3 - SIRT1 co-immunoprecipitation (**Figure 5C-D**) - there was only one threonine residue detected by phosphoproteomics that fell within the amino acid 115 - 464 domain of SIRT1: T344. As T344 is located within the deacetylase domain of SIRT1, post-translational modification of this amino acid residue is likely to affect SIRT1 enzymatic activity.

To determine if MAP4K3 phosphorylation of SIRT1 at T344 might have regulatory significance, we derived a phosphomimetic version of SIRT1 by mutating T344 to an aspartic acid (D) residue. When we transfected MAP4K3 k.o. HEK293A cells with the SIRT1-T344D mutant and tracked mTORC1 activation upon amino acid stimulation in comparison to MAP4K3 k.o. cells expressing empty vector, we observed significant increases in phosphorylation of S6 and 4EBP1 for MAP4K3 k.o. cells expressing the SIRT1-T344D mutant (**Figure 6A-B**), indicative of rescue of mTORC1 activation in cells lacking MAP4K3. We investigated this phenomenon further by measuring the extent of physical interaction between SIRT1 and LKB1 in MAP4K3 k.o. cells, comparing wild-type SIRT1 to SIRT1-T344D. After confirming that both wild-type SIRT1-HA and SIRT1-T344D-HA localized to the nucleus when transfected into MAP4K3 k.o. cells (**Figure S4**), we detected a roughly two-fold increase in the interaction between SIRT1-T344D and LKB1 in MAP4K3 k.o. cells in comparison to the interaction between wild-type SIRT1 and LKB1 (**Figure 6C-D**). Enhanced interaction of SIRT1-T344D with LKB1 is consistent with inhibition of SIRT1 enzymatic deacetylase activity upon T344 phosphorylation, as documented previously (Sasaki *et al*, 2008), and contributes to LKB1 retention in the nucleus, thereby preventing LKB1 activation of AMPK in the cytosol and downstream mTORC1 repression in MAP4K3 k.o. cells. Even though co-transfection, coimmunoprecipitation studies of MAP4K3 and SIRT1 truncation fragments revealed an interaction involving the SIRT1 domain containing T344, it is possible that MAP4K3 phosphorylation of the other two threonine sites (T530 and T719) could have regulatory significance. To directly examine this possibility, we transfected MAP4K3 k.o. HEK293A cells with SIRT1-T530D or SIRT1 T719D, confirmed that both these mutants localized to the nucleus (**Figure S4**), and evaluated mTORC1 activation upon amino acid stimulation in comparison to MAP4K3 k.o. cells expressing empty vector. We found that neither the SIRT1-T530D nor the SIRT1-719D phosphomimetic mutant could rescue mTORC1 activation in MAP4K3 k.o. cells (**Figure S5A-B**), indicating that of the three putative threonine phosphorylation sites, only SIRT1 T344 appears subject to MAP4K3 phosphoregulatory control.

**Figure 6.**
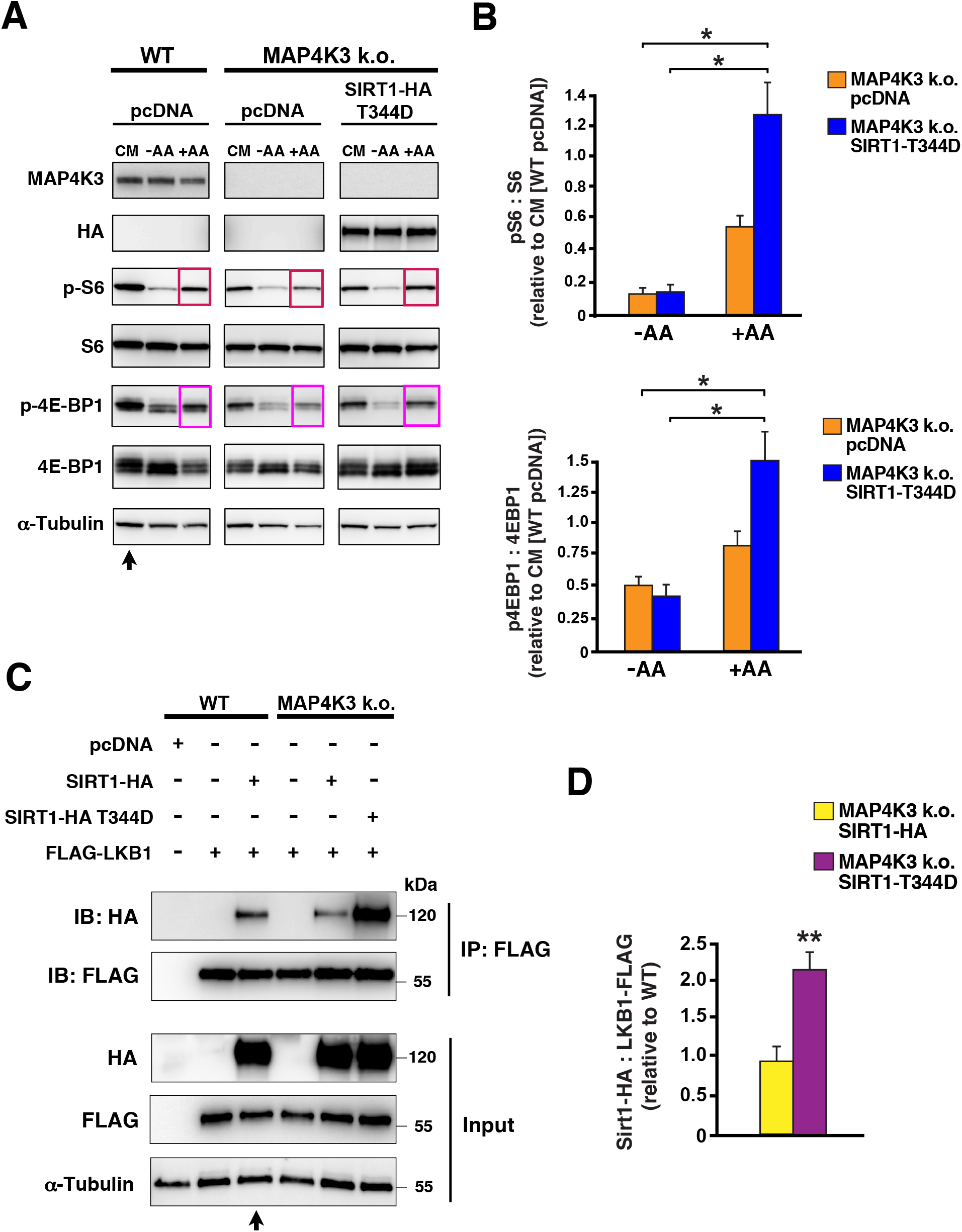
Phosphorylation status of threonine 344 in SIRT1 regulates amino acid dependent activation of mTORC1. **(A)** We transfected MAP4K3 HEK293A k.o. cells with the SIRT1-T344D phosphomimetic mutant or pcDNA empty vector, and maintained the cells under baseline complete media (CM) conditions, or subjected the cells to amino acid starvation (-AA) or amino acid starvation followed by amino acid restimulation (+AA). As a basis of comparison, we transfected WT HEK293A cells with pcDNA empty vector, and maintained them under baseline complete media (CM) conditions, or subjected the cells to amino acid starvation (-AA) or amino acid starvation followed by amino acid restimulation (+AA). We immunoblotted resulting cell protein lysates for MAP4K3, HA-tagged SIRT1, phosphorylated S6, total S6, phosphorylated 4E-BP1, and total 4E-BP1, as indicated. Note rescue of mTORC1 activation in MAP4K3 k.o. cells expressing the SIRT1-T344D mutant. a-tubulin served as the loading control. **(B)** TOP GRAPH: We quantified the levels of phosphorylated S6 and total S6 in **(A)** by densitometry, determined the ratio of phosphorylated S6: total S6, and normalized the results to pcDNA-transfected WT HEK293A cells at baseline (arrow in **(A)**). \*\**P* < 0.01, ANOVA with post-hoc Tukey test. BOTTOM GRAPH: We quantified the levels of phosphorylated 4E-BP1 and total 4E-BP1 in **(A)** by densitometry, determined the ratio of phosphorylated 4E-BP1: total 4E-BP1, and normalized the results to pcDNA-transfected WT HEK293A cells at baseline (arrow in **(A)**). \*\**P* < 0.01, ANOVA with post-hoc Tukey test; n = 3 biological replicates. Error bars = s.e.m. **(C)** We transfected WT HEK293A cells with FLAG-LKB1 alone or in combination with WT SIRT1-HA, or with pcDNA empty vector, and transfected MAP4K3 HEK293A k.o. cells with FLAG-LKB1 alone, FLAG-LKB1 and WT SIRT1-HA, or FLAG-LKB1 and SIRT1-HA-T344D. We then performed co-immunoprecipitation of LKB1 and SIRT1 by FLAG IP, followed by immunoblotting with anti-HA antibody or anti-FLAG antibody. Immunoblotting of protein lysates from input cells is shown below. **(D)** Quantification of SIRT1-LKB1 interaction in **(C)** based upon densitometry analysis performed on WT HEK293A cells co-transfected with FLG-LKB1 and WT-SIRT1-HA, MAP4K3 k.o. cells cotransfected with FLAG-LKB1 and WT SIRT1-HA, and MAP4K3 k.o. cells co-transfected with FLAG-LKB1 and SIRT1-HA-T344D. Results for SIRT1: LKB1 were normalized to WT HEK293A cells cotransfected with FLG-LKB1 and WT-SIRT1-HA (arrow in **(C)**). *P* < 0.01, two-tailed t-test; n = 3 biological replicates. Error bars = s.e.m.

## Discussion

When deciding whether to adopt an anabolic or catabolic state, the cell must integrate numerous inputs reflecting nutrient status and growth potential. The mTORC1 complex sits at the center of this integration process, weighing diverse inputs from multiple signaling pathways. Availability of nitrogen is a very important input; hence, supply of essential amino acids is among the most potent determinants of mTORC1 complex activity and consequently the metabolic state of the cell. For decades, it has been known that MAP4K3 is required for complete and robust activation of the mTORC1 complex (Bryk *et al*., 2010; Findlay *et al*., 2007; Resnik-Docampo & de Celis, 2011), yet the pathway from MAP4K3 to mTORC1 has remained ill-defined. Here we sought to delineate this signaling pathway, after confirming that MAP4K3 is indeed necessary for mTORC1 activation, by studying cell growth and quantifying mTORC1 target phosphorylation in two independently derived stable MAP4K3 k.o. cell lines. We found that MAP4K3 loss-of-function blunted both cell growth and mTORC1 complex activation, and we documented that MAP4K3 activation of mTORC1 operates via suppression of the TSC1/2 complex to de-repress Rheb. When we considered different inputs to the TSC1/2 complex, we found that MAP4K3 represses the LKB1-AMPK axis to drive mTORC1 activation, documenting complete rescue of mTORC1 activation in both MAP4K3/AMPK double k.o. cells and MAP4K3/LKB1 double k.o. cells. To identify the link between MAP4K3 and LKB1, we performed an unbiased interactome screen of MAP4K3, and upon noting evidence for an interaction between MAP4K3 and SIRT1, we confirmed this to be a direct physical interaction. We then performed phosphopeptide and phospho-amino acid analysis on SIRT1 in WT and MAP4K3 k.o. cells, and we examined the physiological relevance of SIRT1 modulation of MAP4K3-dependent activation of mTORC1, focusing on three putative threonine phosphorylation sites. Our results reveal a novel signaling pathway (**Figure 7**), in which MAP4K3 phospho-inhibition of SIRT1 prevents LKB1-AMPK activation of TSC1/2, thereby insuring that Rheb can potently activate mTORC1 at the surface of the lysosome.

**Figure 7.**
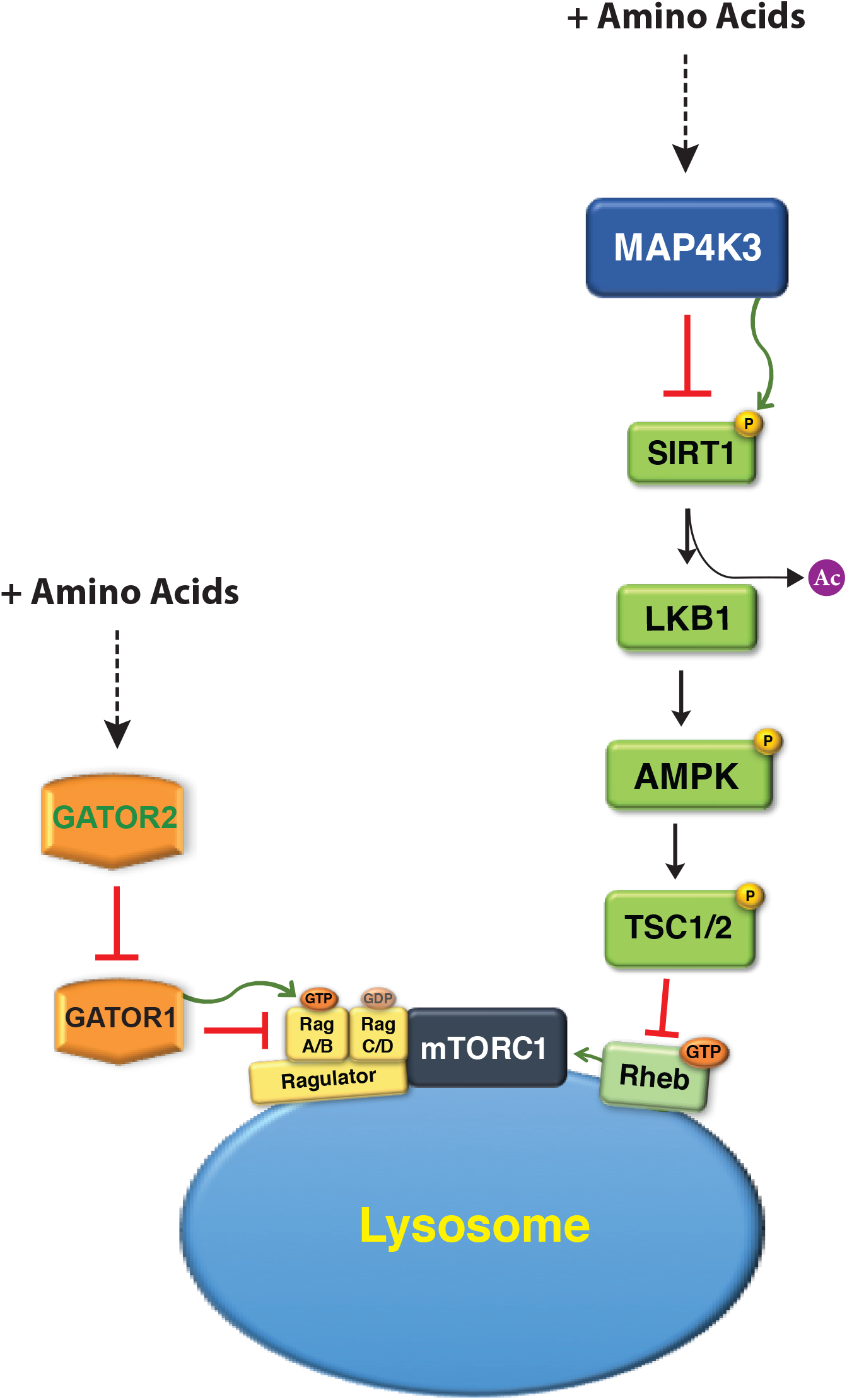
Model for MAP4K3 amino acid dependent activation of the mTORC1 complex. In the absence of amino acids, activated SIRT1 deacetylates LKB1, which phosphorylates AMPK to activate it, resulting in phospho-activation of the TSC1/2 complex. TSC2 is a GTPase activating protein that turns off Rheb by converting GTP to GDP, thereby preventing Rheb activation of the mTORC1 complex. However, when amino acids are plentiful, mTORC1 is recruited to the surface of the lysosome by interaction of the Ragulator complex with the Rag proteins, which are activated by GATOR1. In parallel with subcellular localization of mTORC1 to the lysosome, amino acids activate MAP4K3, which turns off SIRT1 by directly phosphorylating it. With SIRT1 inhibited, LKB1 remains acetylated in the nucleus and cannot activate AMPK. With AMPK inactive, the TSC1/2 complex is repressed, and thus cannot inhibit Rheb. Disinhibited Rheb, which is located on the surface of the lysosome in association with mTORC1, can thus fully activate the mTORC1 complex.

SIRT1 is a NAD^+^-dependent deacetylase, implicated in the regulation of multiple metabolic pathways and shown to promote beneficial caloric-restriction phenotypes of improved health span and in certain instances maximal lifespan (Longo & Kennedy, 2006). In model organisms, the effect of over-expression of Sirtuin orthologues in extending lifespan, though controversial, stand in direct genetic opposition to mTOR orthologues, which upon deletion result in increased lifespan (Houtkooper *et al*, 2012). Here we show that MAP4K3 achieves full activation of mTORC1 by inhibiting SIRT1, which is consistent with a prior study that reported SIRT1 repression of mTORC1 activity in a TSC1/2-dependent manner, postulating an interaction between SIRT1 and TSC2 (Ghosh *et al*, 2010), though a mechanistic explanation for direct SIRT1 regulation of the TSC1/2 complex has not emerged in the decade since publication. Indeed, in almost all cases, the pleiotropic actions of SIRT1 and mTORC1 on metabolic and stress response pathways are diametrically opposed; hence, our discovery of MAP4K3 inhibition of SIRT1-mediated repression of mTORC1 suggests that MAP4K3 is reinforcing the positive regulation of anabolism by activating mTORC1, repressing autophagy, and inhibiting SIRT1. The fact that MAP4K3 knock-out mice display increased lifespan supports this interpretation of previous studies and our current findings (Chuang *et al*, 2019).

The regulation of SIRT1 activity has been the subject of numerous studies, and although the role of post-translational modifications in modulating SIRT1 activity remains unclear, prior investigations have documented numerous SIRT1 phosphorylation sites (Lee *et al*, 2012; Sasaki *et al*., 2008). In agreement with prior studies, our phosphoproteomics analysis identified 17 phosphorylation sites on SIRT1, including T344, T530, and T719. All of these threonine phosphorylation sites have been evaluated in previous studies (Lau *et al*, 2014; Ling *et al*, 2018; Sasaki *et al*., 2008; Shan *et al*, 2017). In 2012, one group first identified SIRT1 T344 as a phosphorylation site for AMPK, and reported that AMPK phosphorylation of SIRT1 T344 yielded inactivation of SIRT1 deacetylation of p53 in liver cancer cells (Sasaki *et al*., 2008). However, subsequent studies focused on the inhibitory interaction of SIRT1 with Deleted in Breast Cancer-1 (DBC1) protein (Kim *et al*, 2008; Zhao *et al*, 2008), and proposed that AMPK and Aurora kinase A may phosphorylate SIRT1 at T344 to release it from interaction with DBCl, thereby promoting SIRT1 activation (Lau *et al*., 2014; Ling *et al*., 2018). Unlike the initial 2012 report, these latter studies, however, did not directly assay the deacetylase activity of SIRT1 T344 phospho-mutants for their proposed targets, leaving open the possibility that an alternate phosphorylation or post-translational modification mediates the observed biological effects. Our results by no means preclude the existence of additional regulatory phosphorylation sites on SIRT1; indeed, we predict that other post-translational modifications or modes of regulation may determine SIRT1 enzymatic activity and function. Furthermore, while SIRT1 deacetylation of LKB1 has been shown to promote LKB1 activation (Lan *et al*., 2008), LKB1 is in complex with the pseudokinase STRAD– and scaffold protein MO25 (Kullmann & Krahn, 2018), suggesting that these factors may coordinate LKB1 regulation. Whether and how SIRT1 activity is controlled by phosphorylation at other sites in addition to T344 to regulate mTORC1 activation via the LKB1-AMPK pathway and how complete LKB1 activation is fully achieved should thus be the focus of future studies.

Our findings indicate that MAP4K3 is a central point of integration for nutrient sensing regulation in the cell. We previously reported that MAP4K3 supersedes mTORC1 in the regulation of TFEB-dependent autophagy activation (Hsu *et al*., 2018). Here we have shown that MAP4K3 is required for amino acid dependent activation of mTORC1, and through its regulation of SIRT1 and AMPK, MAP4K3 is serving as a point of cross-talk between mTORC1 and other well-recognized master regulators. While we do not yet know how MAP4K3 senses amino acids to become activated, we found that MAP4K3 is required for full mTORC1 activation by leucine and arginine. As leucyl-tRNA synthetase 1 (LARS) only activates the mTORC1 complex when both leucine and glucose are in abundant supply (Yoon *et al*, 2020), some amino acid sensing regulators appear to be subject to modulation by glucose levels as well as amino acid supply. While MAP4K3 robustly mediates amino acid dependent activation of mTORC1, MAP4K3 may not be responsive to glucose levels, as MAP4K3 loss-of-function did not affect mTORC1 activation in the presence of glucose and did not alter mTORC1 activity in response to glucose in the presence or absence of AMPK. However, whether other metabolic sensors responding to glucose or to other nutrient levels interact with MAP4K3 to regulate its activity via cross-talk deserves further consideration in different cell types and under different circumstances.

### Inhibition of MAP4K3 as a potential therapeutic strategy

The physiological relevance of MAP4K3 function for various disease processes has been the subject of prior investigation, and one potentially important role for MAP4K3 is in the regulation of immune system function, where MAP4K3 has been shown to directly activate Protein Kinase C-θ in T cells (Chuang *et al*, 2011). Furthermore, MAP4K3 loss-of-function renders knock-out mice resistant to experimental autoimmune encephalitis, and human patients with systemic lupus erythematosus display increased expression of MAP4K3 accompanied by hyperactivation of Protein Kinase C-θ (Chuang *et al*., 2019; Chuang *et al*., 2011). In addition to potential modulation of MAP4K3 as a treatment for autoimmune disease, there are many disorders believed to involve over-activation of mTORC1, including various cancers and certain neurological diseases (Lipton & Sahin, 2014; Zou *et al*, 2020). The problem with deploying drug inhibitors of mTOR in human patients has been the occurrence of side effects and adverse events (Pallet & Legendre, 2013), limiting the dosages of rapamycin analogues, or so-called “rapalogues”. Though MAP4K3 is a potent input to mTORC1, its effect upon mTORC1 in cellular physiology is likely balanced by inputs from other master regulators, suggesting that MAP4K3 inhibition may not produce the extensive, deleterious physiological effects observed upon mTOR inhibition. Indeed, as MAP4K3 loss-of-function does not result in disease phenotypes in knock-out mice (Chuang *et al*., 2019), it is possible that MAP4K3 inhibitors will be better tolerated than rapalogues, and thus may yield a novel class of drugs for use in human patients afflicted with diseases stemming from mTORC1 hyperactivation.

## Materials and Methods

### Cell culture and amino acid treatments

HEK293A cells (Thermo Fisher #R70507) were grown in and maintained in complete media (high glucose DMEM containing 10% FBS (Gibco)). Derivation and characterization of two independently generated MAP4K3 knock-out cell lines from the HEK293A cells were previously described (Hsu *et al*., 2018). For amino acid deprivation (-AA), cells were washed once with Earl’s Balance Salt Solution (EBSS) and maintained in amino acid-free media (EBSS containing 10% dialyzed FBS (Gibco #A3382001)) for time intervals indicated. Restimulation was performed by replacing amino acid-free media with low glucose DMEM (~1 g/L D-glucose) containing 10% dialyzed FBS for indicated time intervals. Transfections were performed using Lipofectamine 2000, according to the manufacturer’s instructions (Invitrogen). For qRT-PCR experiments, transfection was performed with 2.4 mg of DNA per 10 cm^2^ of cells. For immunofluorescence experiments, transfection was performed with 0.08 mg of DNA per 0.7 cm^2^ of cells. After 6 hrs, the media was replaced. Complete media consisting of high glucose was prepared as follows: D-glucose 4.5 g/L, L-glutamine 584.0 mg/L.

### Generation of MAP4K3 double knock-out cell lines

The 20 nucleotide guide sequences targeting human AMPK a1 subunit and LKB1 were designed using the CRISPR design tool at http://crispr.mit.edu/ (Hsu *et al*, 2013) and cloned into a bicistronic expression vector (pX330) containing human codon-optimized Cas9 and RNA components (Addgene, #42230). The guide sequences targeting the AMPKα1 gene *(PRKAA1)* in exon 1 were as follows:

Site 1: 5’-CACCGGAAGATCGGCCACTACATTC-3’; 5’-AAACGAATGTAGTGGCCGATCTTCC-3’
Site 2: 5’-CACCGGAAGATCGGACACTACGTGC-3’; 5’-AAACGCACGTAGTGTCCGATCTTCC-3’

The guide sequences targeting the LKB1 gene in exon 1 were as follows:

Site 1: 5’-CACCGAGCTTGGCCCGCTTGCGGCG-3’; 5’-AAACCGCCGCAAGCGGGCCAAGCTC-3’
Site 2: 5’-CACCGGTTGCGAAGGATCCCCAACG-3’; 5’-AAACCGTTGGGGATCCTTCGCAACC-3’

Single guide RNAs (sgRNAs) in the pX330 vector (1 μg) were mixed with EGFP (0.1 μg; Clontech) and co-transfected into MAP4K3 k.o. HEK293A cells (line 1) using Lipofectamine 2000 (Life Technologies) according to manufacturer’s instructions. 24 hrs post transfection, the cells were trypsinized, washed with PBS, and re-suspended in fluorescence-activated cell sorting (FACs) buffer (PBS, 5 mM EDTA, 2% FBS and Pen/Strep). GFP positive cells were single cell sorted by FACs (BDInflux) into 96-well plate format into DMEM containing 20% FBS and 50 μg ml/L penicillin /streptomycin. Single clones were expanded, and we screened for loss of the AMPK α1 subunit or LKB1 protein by immunoblotting. Genomic DNA (gDNA) was purified from clones using the DNeasy Blood & Tissue Kit (QIAGEN, #69504), and the region surrounding the protospacer adjacent motif (PAM) was amplified with Phusion^®^ High-Fidelity DNA Polymerase (New England Biolabs, #M0530). PCR products were purified using the QIAquick PCR Purification Kit (QIAGEN, #28104) and cloned using the TOPO^®^ TA Cloning (ThermoFisher, #K457502). To determine the specific mutations for individual alleles, at least 10 different bacterial colonies were expanded and the plasmid DNA purified and sequenced.

### Cell lysis and immunoprecipitation

Cells were rinsed twice with ice-cold PBS and lysed in ice-cold lysis buffer (25mM HEPES-KOH pH 7.4, 150mM NaCL, 5mM EDTA, 1% Triton X-10040 mM, one tablet of EDTA-free protease inhibitors (Roche, #11873580001) per 10 mL of lysis buffer, and one tablet of PhosStop phosphatase inhibitor (Roche, #4906845001), as necessary. The soluble fractions from cell lysates were isolated by centrifugation at 8,000 rpm for 10 min in a microfuge. For immunoprecipitations, primary antibodies were incubated with Dynabeads^®^ (Invitrogen) overnight, then washed with sterile PBS. Antibodies bound to Dynabeads were then incubated with lysates with rotation for 2 hrs at 4°C. Immunoprecipitates were washed three times with lysis buffer. Immunoprecipitated proteins were denatured by the addition of 20 μl of sample buffer and boiling for 10 min at 70°C, resolved by SDS-PAGE, and analyzed via Western blot analysis.

### Western blot analysis

Protein quantification was performed with Pierce Rapid Gold BCA (ThermoFisher, #A53225) and proteins were denatured with LDS sample buffer and boiled for 10 min at 70 °C. 30-35 μg of protein were loaded into each well, resolved by SDS-polyacrylamide gel electrophoresis (PAGE), and analyzed by immuoblot analysis with the indicated antibodies (-see Table below; dilutions available upon request). Species-specific secondary antibodies were goat anti-rabbit IgG-HRP (Santa Cruz, #sc-2004) or goat anti-mouse IgG-HRP (Santa Cruz, #sc-2005), diluted 1/10,000 in 5% PBS-T milk and incubated for 1 hr at RT. Chemiluminescent signal detection was captured with Pierce ECL Plus Western Blotting Substrate (ThermoFisher, #321-32), and autoradiography film, using standard techniques. Levels of total and phosphorylated protein were analyzed on separate gels and normalized to β-actin or α-Tubulin (loading control). Band intensities were determined using densitometry analysis on ImageJ software (NIH).

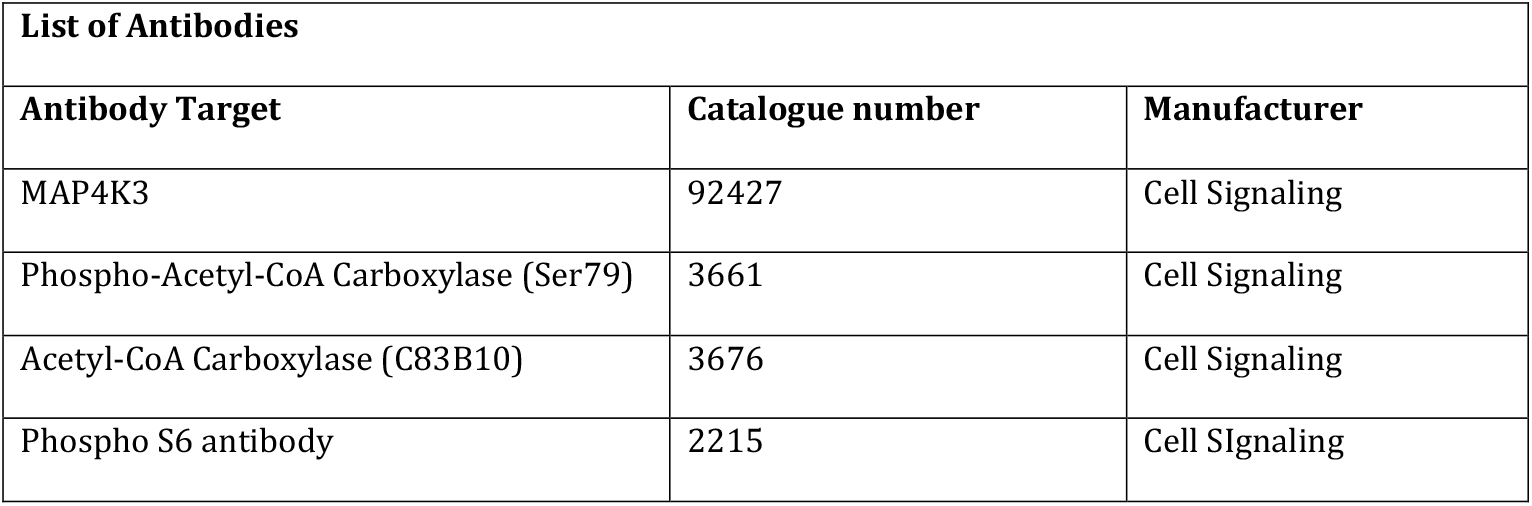
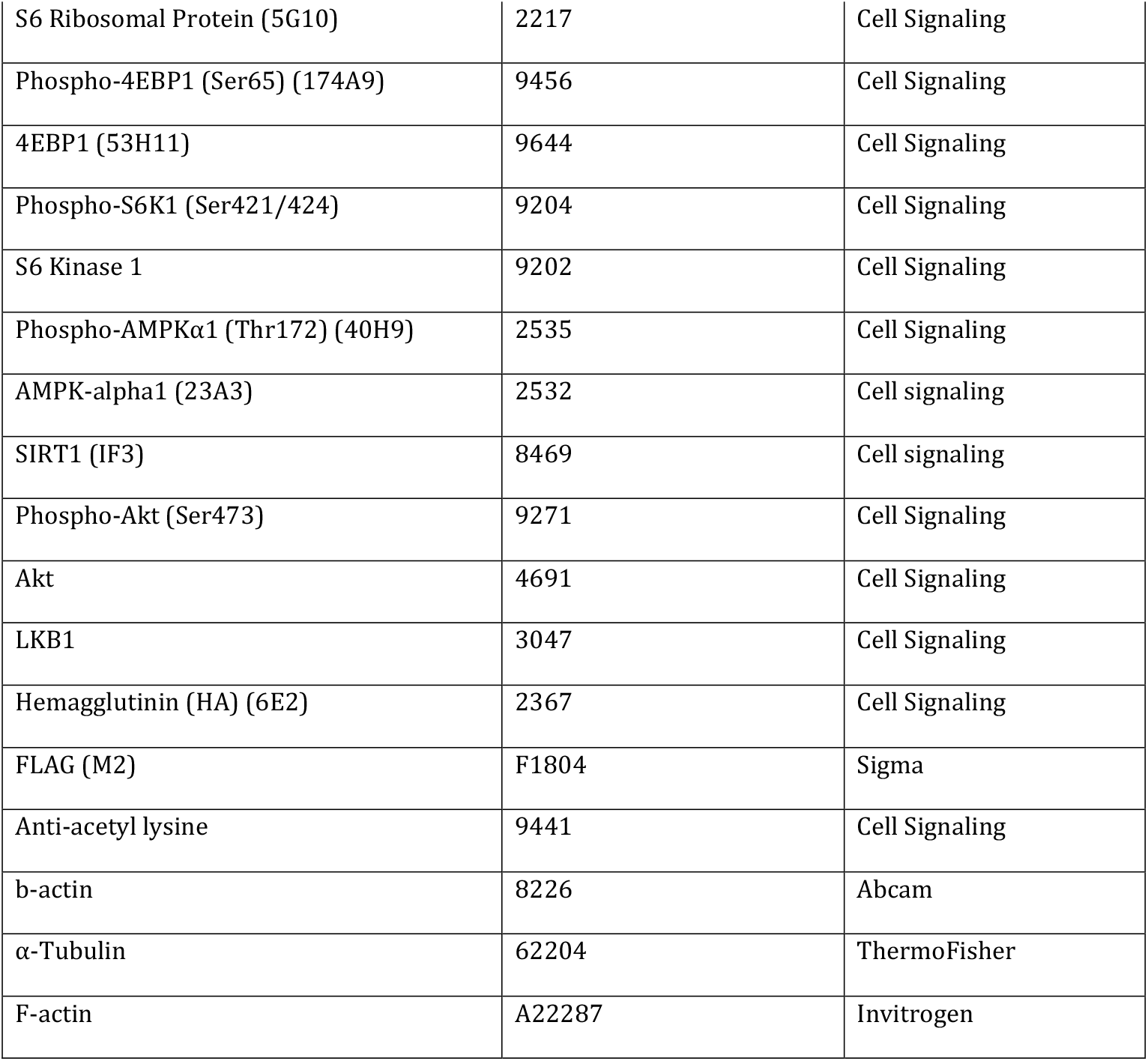

| **List of Antibodies** |  |  |
| --- | --- | --- |
| **Antibody Target** | **Catalogue number** | **Manufacturer** |
| MAP4K3 | 92427 | Cell Signaling |
| Phospho-Acetyl-CoA Carboxylase (Ser79) | 3661 | Cell Signaling |
| Acetyl-CoA Carboxylase (C83B10) | 3676 | Cell Signaling |
| Phospho S6 antibody | 2215 | Cell Signaling |
| S6 Ribosomal Protein (5G10) | 2217 | Cell Signaling |
| Phospho-4EBP1 (Ser65) (174A9) | 9456 | Cell Signaling |
| 4EBP1 (53H11) | 9644 | Cell Signaling |
| Phospho-S6K1 (Ser421/424) | 9204 | Cell Signaling |
| S6 Kinase 1 | 9202 | Cell Signaling |
| Phospho-AMPKα1 (Thr172) (40H9) | 2535 | Cell Signaling |
| AMPK-alpha1 (23A3) | 2532 | Cell signaling |
| SIRT1 (IF3) | 8469 | Cell signaling |
| Phospho-Akt (Ser473) | 9271 | Cell Signaling |
| Akt | 4691 | Cell Signaling |
| LKB1 | 3047 | Cell Signaling |
| Hemagglutinin (HA) (6E2) | 2367 | Cell Signaling |
| FLAG (M2) | F1804 | Sigma |
| Anti-acetyl lysine | 9441 | Cell Signaling |
| b-actin | 8226 | Abcam |
| α-Tubulin | 62204 | ThermoFisher |
| F-actin | A22287 | Invitrogen |

### Immunocytochemistry

Cells were seeded in CC2-coated 8-chamber slides (ThermoFisher, #154941) two days prior to experimentation and transfected as indicated. PBS-MC (1mM MgCl_2_, 0.1mM CaCl_2_, in PBS) was used for all washes and as a diluent for all solutions. Cells were fixed with 4% paraformaldehyde in PBS-MC for 12 min, then washed 3 times. Then 0.05% Triton-X in PBS-MC was used to permeabilize the cells for 5 min, followed by 2 washes in PBS-MC. Primary antibodies used are given above, all diluted in 5% normal goat serum in PBS-MC. Cells were incubated in primary antibodies for 2 hrs at RT, followed by 4 washes in PBS-MC. Cells were incubated in secondary antibodies (Alexa Fluor, ThermoFisher) in 5% NGS in PBS-MC for 1 hr at RT. Cells were washed 4 times in PBS-MC, then mounted with Prolong gold antifade reagent with DAPI (ThermoFisher, #P-36931) or Hoechst. Images were captured with a Zeiss LSM 780 confocal microscopy or Zeiss LSM 880 airy scan, and analyzed with Zen 2011 LSM 780 software and Image J.

### Phosphopeptide mapping and phosphoamino acid analysis

SIRT1-HA was first overexpressed in WT HEK293A and MAP4K3 k.o. cells using transfection with Lipofectamine 2000. After transfection, the cell media was changed to DMEM-PO4 with added P^32^ orthophosphate and incubated for 16 hrs. Then, cells were starved or stimulated with amino acids for 30 min. Lysates were prepared, immunoprecipitated with HA antibody for SIRT1 protein, and analyzed by SDS-PAGE, followed by autoradiography of P^32^ incorporation into SIRT1 with a phosphorimager. For phosphopeptide mapping, P^32^-labeled SIRT1 was extracted from the dried gel and precipitated with TCA. The precipitated protein was oxidized, digested with glutamyl endopeptidase, lyophilized, and the tryptic peptide mix spotted onto a thin layer cellulose (TLC) plate (van der Geer & Hunter, 1994). The peptides were then resolved by electrophoresis and chromatography in two dimensions on TLC plates and visualized by autoradiography (van der Geer & Hunter, 1994). For Phospho-amino acid analysis, 50 cpm of purified P^32^-labeled HA-SIRT1 protein was hydrolyzed by incubation for 60 min at 110°C in 30 μl of 6N HCl. The sample was then mixed with stainable phosphoserine, phosphothreonine and phosphotyrosine standards and resolved in two dimensions on TLC plates by electrophoresis. The phospho-amino acid composition was determined by matching the resultant spots on the autoradiograph with the positions of the added ninhydrin-stained standards on the TLC plate.

### Mass spectrometry

FLAG epitope-tagged MAP4K3 constructs were transfected into HEK293T cells and immunoprecipitated as described above. The sample was then run on an 8-16% gel, and analyzed as previously described (Freibaum *et al*, 2010).

For the phosphoproteomics, we isolated 12 SIRT1 gel bands (3 replicates each of WT - amino acids, WT + amino acids, MAP4K3 k.o - amino acids, and MAP4K3 k.o. + amino acids). All samples were then reduced for 30 min at 80°C with 10 mM dithiolthreitol and alkylated for 30 min at RT with 25 mM iodoacetamide. Proteins were digested with 10ng/ul of sequencing grade trypsin (Promega) overnight at 37C. After digestion, an extraction solution of 1% formic acid/2% acetonitrile was added and bands were incubated for 30 min at RT. Next, additional peptides were recovered by adding neat acetonitrile to shrink the gel pieces after removing the supernatant. The acetonitrile was removed, combining it with the previous supernatant, and samples were dried. Samples were resuspended in 20 μL 1%TFA/2% acetonitrile containing 12.5 fmol/μL yeast alcohol dehydrogenase (ADH_YEAST) as well as pre-digested bovine alpha casein at 1 or 2 pmol. From each sample, 3 μL was removed to create a QC Pool sample which was run periodically throughout the acquisition period. Quantitative LC-MS/MS was performed on 3 μL of each sample, using a nanoAcquity UPLC system (Waters Corp) coupled to a Thermo Orbitrap Fusion Lumos high resolution accurate mass tandem mass spectrometer (Thermo) via a nanoelectrospray ionization source. Briefly, the sample was first trapped on a Symmetry C18 20 mm × 180 μm trapping column (5 μl/min at 99.9/0.1 v/v water/acetonitrile), after which the analytical separation was performed using a 1.8 μm Acquity HSS T3 C18 75 μm × 250 mm column (Waters Corp.) with a 60-min linear gradient of 3 to 30% acetonitrile with 0.1% formic acid at a flow rate of 400 nanoliters/minute (nL/min) with a column temperature of 55C. Data collection on the Fusion Lumos mass spectrometer was performed in a data-dependent acquisition (DDA) mode of acquisition with a r=120,000 (@ m/z 200) full MS scan from m/z 375 - 1500 with a target AGC value of 4e5 ions. MS/MS scans were acquired at Rapid scan rate (Ion Trap) with an AGC target of 1e4 ions and a max injection time of 100 ms. The total cycle time for MS and MS/MS scans was 2 sec. A 20 sec dynamic exclusion was employed to increase depth of coverage. The total analysis cycle time for each sample injection was ~1.5 hours. A total of 16 UPLC-MS/MS analyses (excluding conditioning runs, but including 4 replicate SPQC injections) were performed; the SPQC pool containing an equal mixture of each sample was analyzed after every 4 samples throughout the entire sample set (4 times total). Resultant data was imported into Proteome Discoverer 2.4 (Thermo Scientific Inc.) and all LC-MS/MS runs were aligned based on the accurate mass and retention time of detected ions (“features”) which contained MS/MS spectra using Minora Feature Detector algorithm in Proteome Discoverer. Relative peptide abundance was calculated based upon area-under-the-curve (AUC) of the selected ion chromatograms of the aligned features across all runs. Peptides were annotated at a maximum 1% peptide spectral match (PSM) false discovery rate. Missing values were imputed in the following manner. If less than half of the values are missing in a treatment group, values are imputed with an intensity derived from a normal distribution defined by measured values within the same intensity range (20 bins). If greater than half values are missing for a peptide in a group and a peptide intensity is >5e6, then it was concluded that peptide was misaligned and its measured intensity is set to 0. All remaining missing values are imputed with the lowest 5% of all detected values. The following analyses are based on these imputed values. The MS/MS data was searched against the SwissProt *H. sapiens* database (downloaded in Nov 2019) and an equal number of reversed-sequence “decoys” for false discovery rate determination. Mascot Distiller and Mascot Server (v 2.5, Matrix Sciences) were utilized to produce fragment ion spectra and to perform the database searches.

### Growth assay

We seeded the indicated cell lines at either 50,000 cells per well, 25,000 cells per well or 12,500 cells per well in 24 well plates for evaluation of cell growth at 24 hrs, 48 hrs, or 72 hrs, respectively, post-seeding. Cell number was evaluated using the Cell Count and Viability Assay on the Nucleocounter NC-3000 (Chemometic). Cell growth was calculated by determining the fold change of cell number at seeding to time analysis.

### Statistical Analysis

All data were prepared for analysis with standard spread sheet software (Microsoft Excel). Statistical analysis was done using Microsoft Excel, Prism 5.0 (Graph Pad), the VassarStats website (http://faculty.vassar.edu/lowry/VassarStats.html), or One-Way ANOVA calculator website (https://astatsa.com/OneWay_Anova_with_TukeyHSD/). For ANOVA, if statistical significance was achieved (*P* < 0.05), we performed post hoc analysis to account for multiple comparisons. All t-tests were two-tailed. The level of significance (alpha) was always set at 0.05.

## Acknowledgments

This work was supported by grants from N.I.H. (R01 AG033082 and R35 NS122140 to A.R.L.S., R01 CA014915, CA080100, and CA082683 to T.H., and T32 GM008666 and AG000216 to C.L.H.), and by the Waitt Advanced Biophotonics Core Facility of the Salk Institute with funding from the N.I.H. (NCI P30 CA014195 and NINDS P30 NS072031) and from the Waitt Foundation. M.R.B. was supported by a Graduate Research Fellowship from the National Science Foundation. K.O. was supported by a JSPS Overseas Research Fellowship from the Japan Society for the Promotion of Science. J.P.T. is supported by the HHMI. T.H. is an American Cancer Society Professor and holds the Renato Dulbecco Chair in Cancer Research.

## Author Contributions

A. R.L.S., M.R.B., C.L.H., and K.O. provided the conceptual framework for the study. M.R.B., C.L.H., K.O., B.L.S., J.P.T., T.H. and A.R.L.S. designed the experiments. M.R.B., C.L.H., K.O., E.L., J.M., B.W., B. L.S., J.P.T., T.H. and A.R.L.S. performed the experiments. A.R.L.S., M.R.B., C.L.H., and K.O. wrote the manuscript.

## Declarations

The authors have nothing to declare.

---

**Supplementary Table 1.**
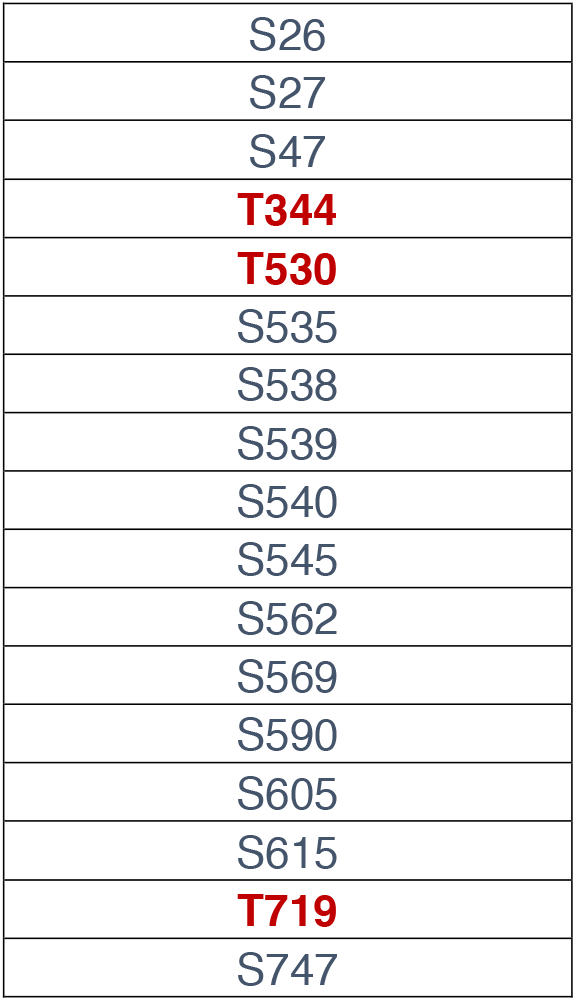
Sirtuin-1 phosphorylation sites detected by mass spectrometry.

|  |
| --- |
| S26 |
| S27 |
| S47 |
| **T344** |
| **T530** |
| S535 |
| S538 |
| S539 |
| S540 |
| S545 |
| S562 |
| S569 |
| S590 |
| S605 |
| S615 |
| **T719** |
| S747 |

## Supplementary Figure Legends

**Figure 1.**
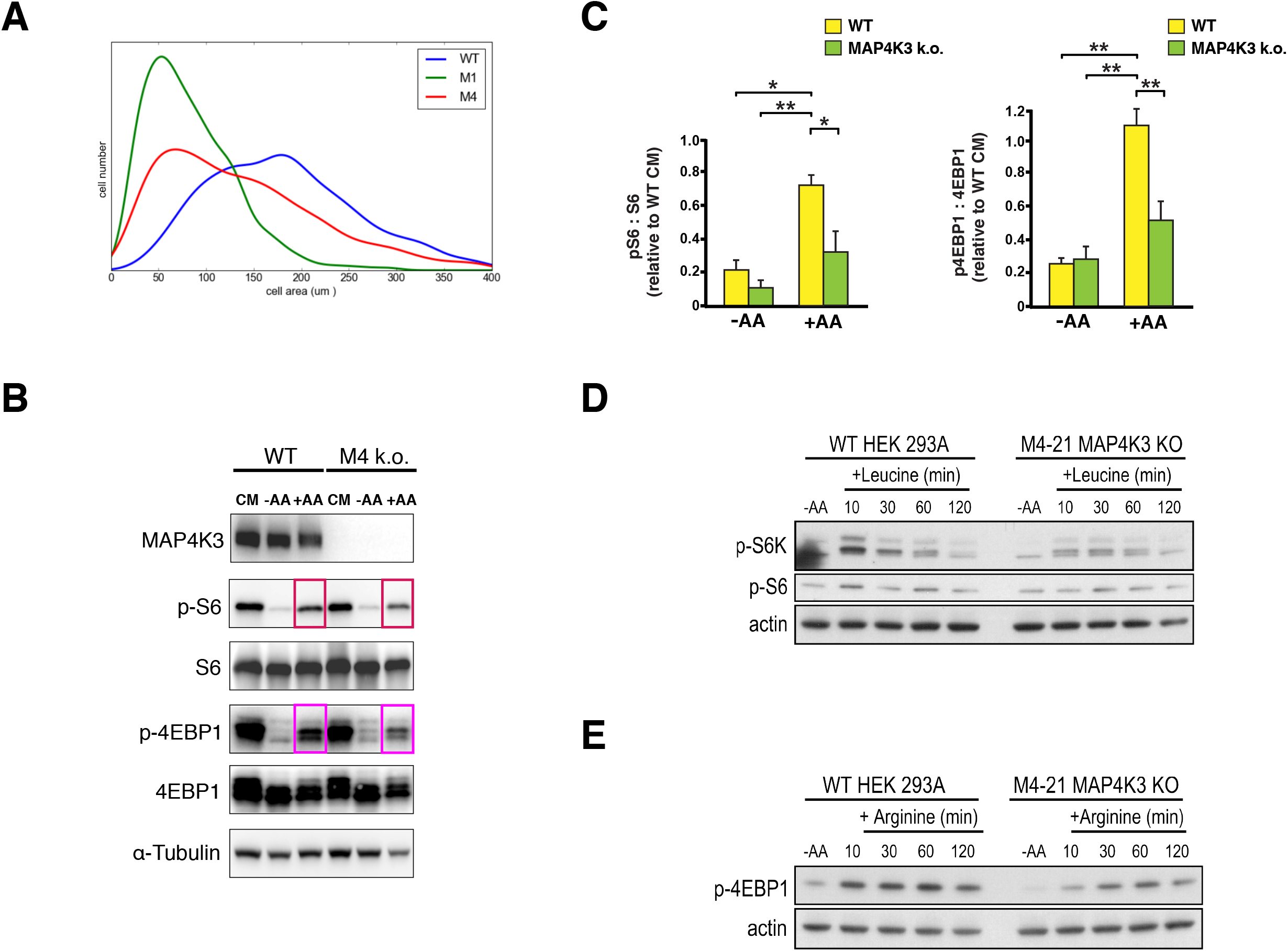
MAP4K3 mediates amino acid-dependent mTORC1 activation and cell proliferation. **(A)** Kernel density plot of a wild-type HEK293A cell line (WT) and two independently derived lines of MAP4K3 knock-out cells (M1, M4) cultured in complete media after 72 hrs. **(B)** WT and MAP4K3 k.o. cells were grown in complete media (CM), starved of amino acids for 60 min (-AA), or starved of amino acids for 60 min and then re-stimulated with amino acids for 20 min (+AA). We immunoblotted resulting cell protein lysates for MAP4K3, phosphorylated S6, total S6, phosphorylated 4E-BP1, and total 4E-BP1, as indicated. Note reduced phospho-activation of mTORC1 targets in MAP4K3 k.o. cells compared to WT cells; a-tubulin served as loading control. **(C)** LEFT: We quantified levels of phosphorylated S6 and total S6 in **(B)** by densitometry, determined the ratio of phosphorylated S6: total S6, and normalized the results to pcDNA-transfected WT HEK293A cells at baseline. \**P* < 0.05, \*\**P* < 0.01; ANOVA with post-hoc Tukey test. RIGHT: We quantified levels of phosphorylated 4E-BP1 and total 4E-BP1 in **(B)** by densitometry, determined the ratio of phosphorylated 4E-BP1: total 4E-BP1, and normalized the results to pcDNA-transfected WT HEK293A cells at baseline. \*\**P* < 0.01, ANOVA with post-hoc Tukey test. n = 3 biological replicates. Error bars = s.e.m. **(D)** WT and MAP4K3 k.o. cells were starved for 120 min, and then re-stimulated with leucine for 10 min, 30 min, 60 min, or 120 min. We immunoblotted the resulting cell protein lysates for phosphorylated S6 kinase 1 and phosphorylated S6. β-actin served as the loading control. **(E)** WT and MAP4K3 k.o. cells were starved for 120 min, and then re-stimulated with arginine for 10 min, 30 min, 60 min, or 120 min. We immunoblotted the resulting cell protein lysates for phosphorylated 4E-BP 1. β-actin served as the loading control.

**Figure 2.**
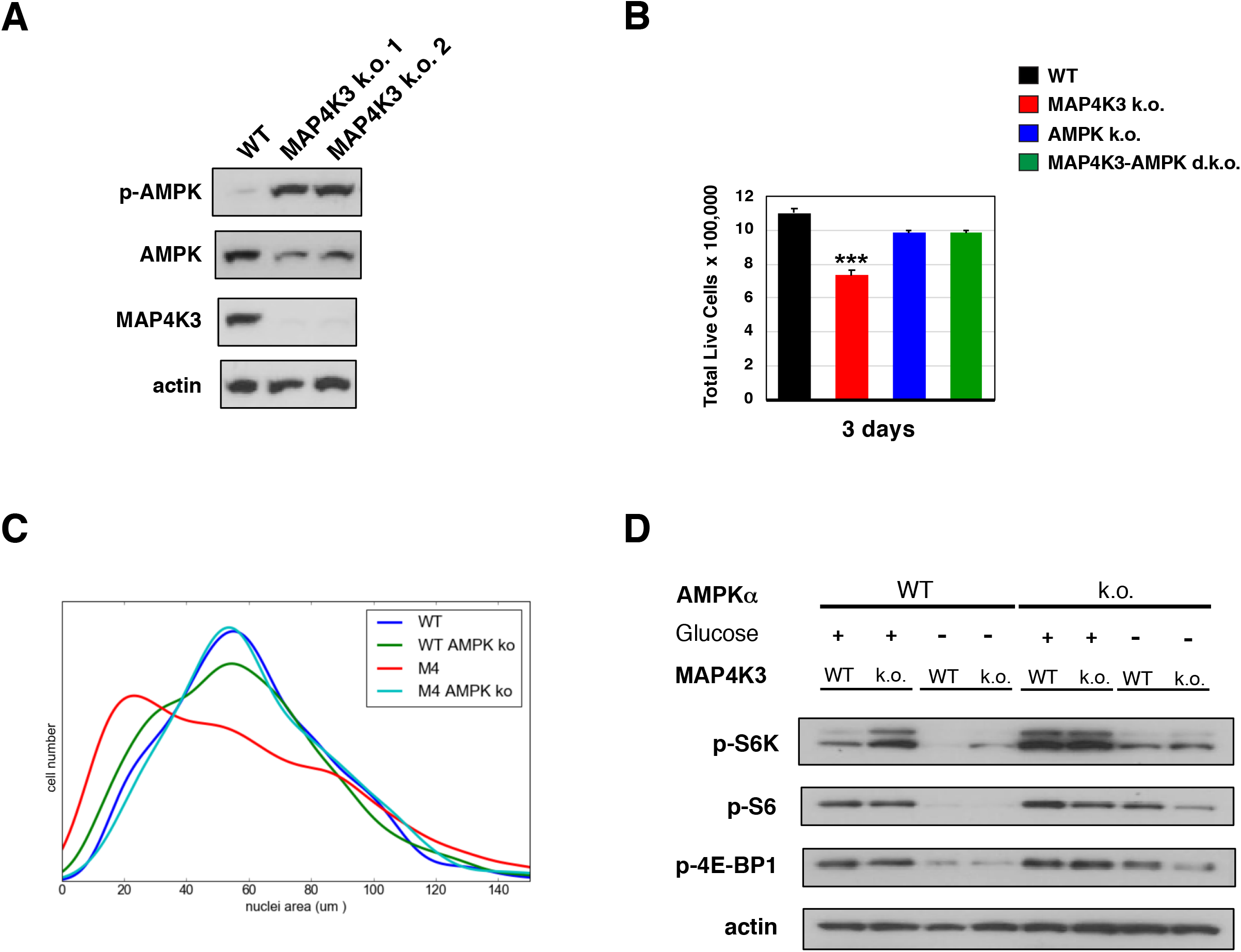
MAP4K3 regulates AMPK to dictate mTORC1 activation in response to amino acids. **(A)** WT cells and two independently derived clonal lines of MAP4K3 k.o. cells were starved of amino acids for 3 hrs and then re-stimulated with amino acids for 30 min. We immunoblotted resulting cell protein lysates for phosphorylated AMPK α1 subunit, total AMPK α1 subunit, and MAP4K3, as indicated. β-actin served as the loading control, and we did not detect any changes in adenine nucleotide levels between WT and MAP4K3 k.o. cells. **(B)** We cultured a WT HEK293A cell line, MAP4K3 k.o. cell line, AMPK α1 k.o. cell line, and MAPK3/AMPK α1 double k.o. cell line in complete media over 72 hrs. Here we see quantification of cell numbers at the end of the 72 hr interval. \*\*\**P* < 0.001; ANOVA with post-hoc Tukey test. n = 3 technical replicates. Error bars = s.e.m. **(C)** Kernel density plot of a WT HEK293A cell line, MAP4K3 k.o. cell line, AMPK α1 k.o. cell line, and MAPK3/AMPK α1 double k.o. cell line cultured in complete media after 72 hrs. **(D)** WT cells, MAP4K3 k.o. cells, AMPK α1 k.o. cells, and MAPK3/AMPK α1 double k.o. cells were cultured in complete media in the presence or absence of glucose for 3 hrs. We immunoblotted resulting cell protein lysates for phosphorylated S6 kinase 1, phosphorylated S6, and phosphorylated 4E-BP1, as indicated. β-actin served as the loading control.

**Figure 3.**
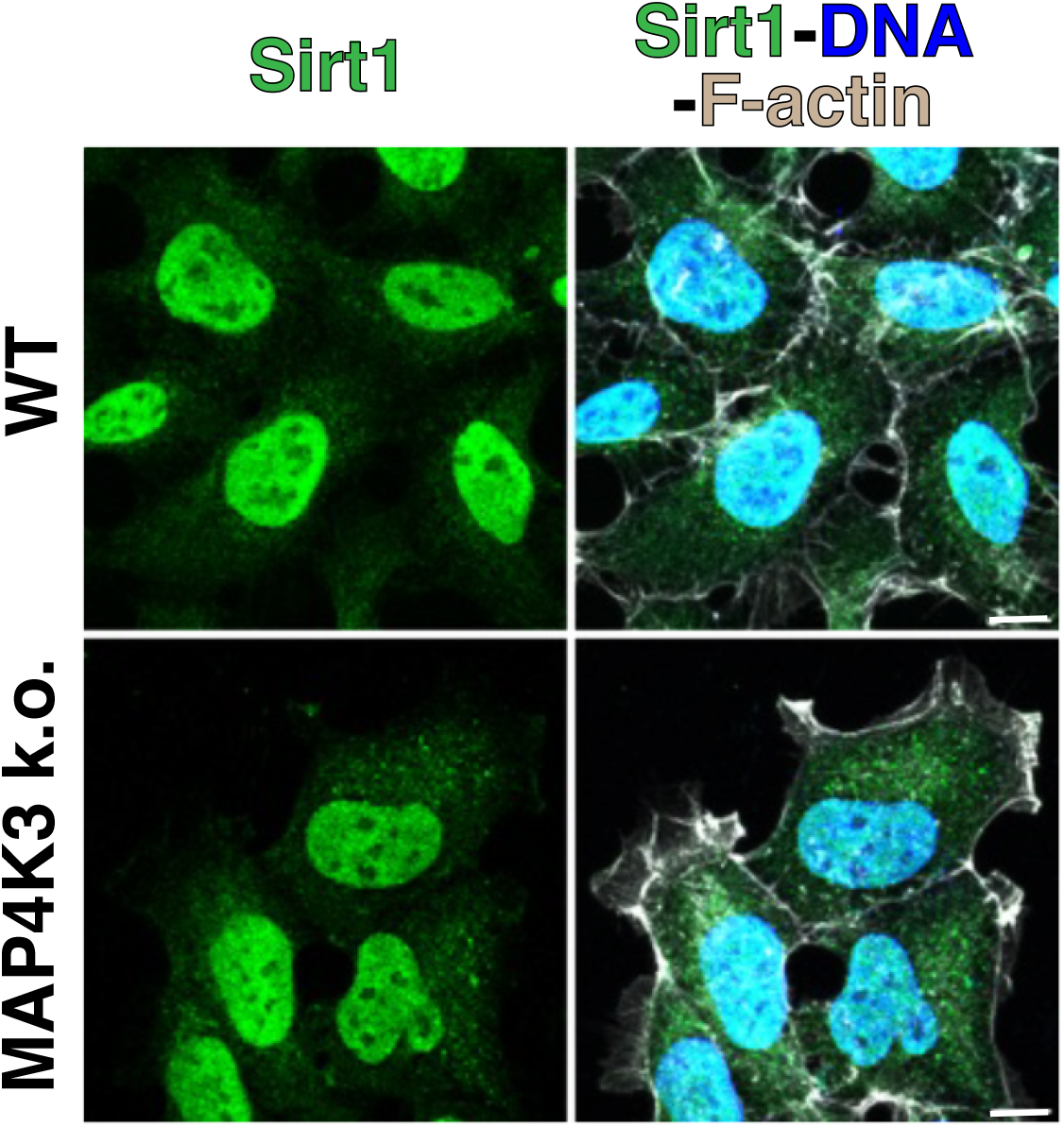
SIRT1-HA localizes to the nucleus in WT and MAP4K3 k.o. HEK293A cells. We transfected a WT HEK293A cell line or a MAP4K3 k.o. cell line with a WT SIRT1-HA expression vector for 24 hrs, immunostained with anti-HA antibody (green) and anti-F-actin antibody (gray), and counterstained with DAPI (blue). Note nearly complete nuclear localization of SIRT1-HA in both WT and MAP4K3 k.o. cells. Scale bars = 10 μm

**Figure 4.**
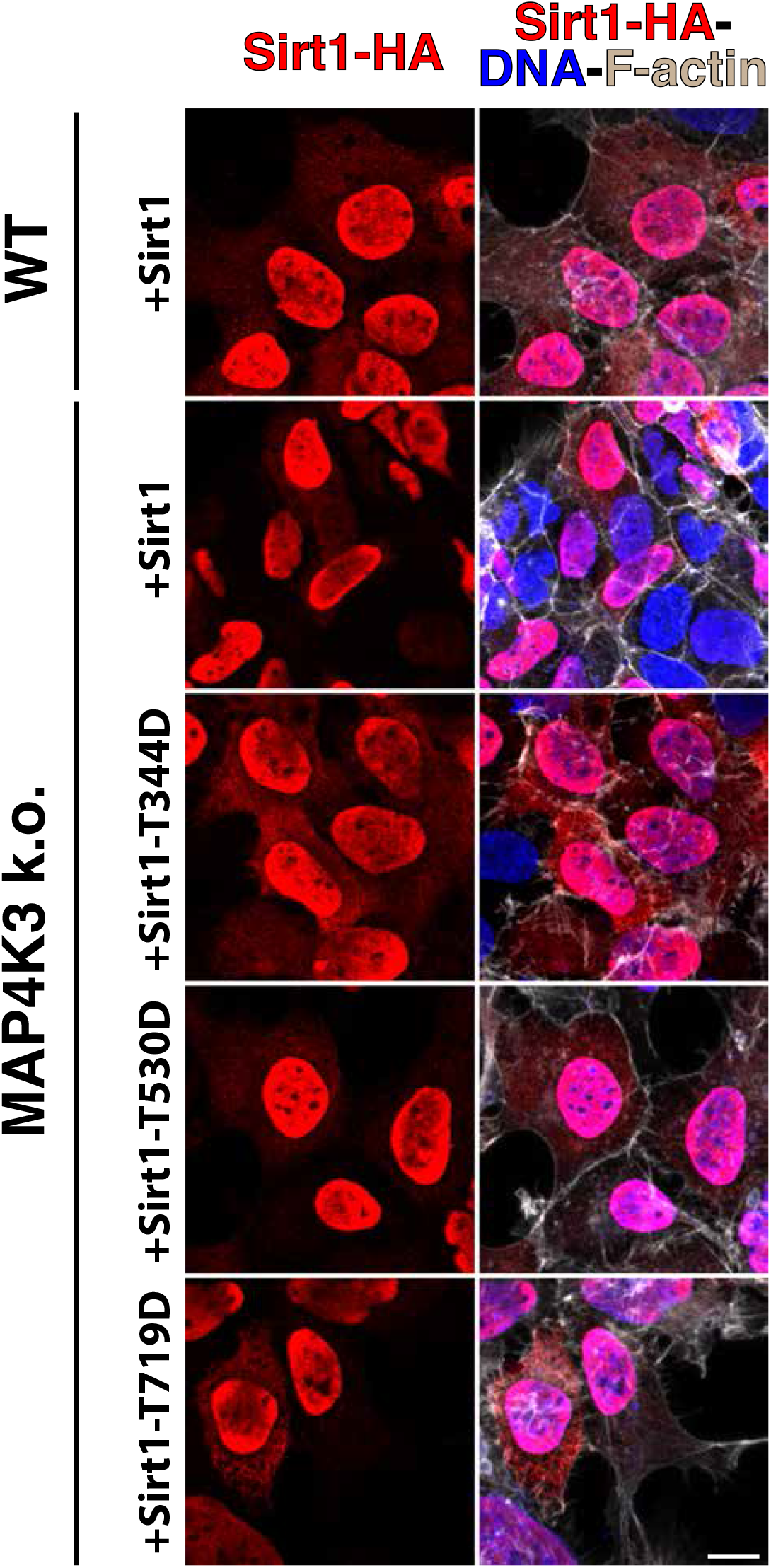
SIRT1 phosphomimetic mutants localize to the nucleus in MAP4K3 k.o. cells. We transfected WT HEK293 cells with a WT SIRT1-HA expression vector for 24 hrs, or transfected a MAP4K3 k.o. cell line with either WT SIRT1-HA, SIRT1-T344D-HA, SIRT1-T530D-HA, or SIRT1-T719D for 24 hrs, immunostained with anti-HA antibody (red) and anti-F-actin antibody (gray), and counterstained with DAPI (blue). Note nearly complete nuclear localization of SIRT1-HA in both WT and MAP4K3 k.o. cells, as well as nearly complete nuclear localization of the various SIRT1 threonine phosphomimetic mutants in MAP4K3 k.o. cells. Scale bars = 10 μm

**Figure 5.**
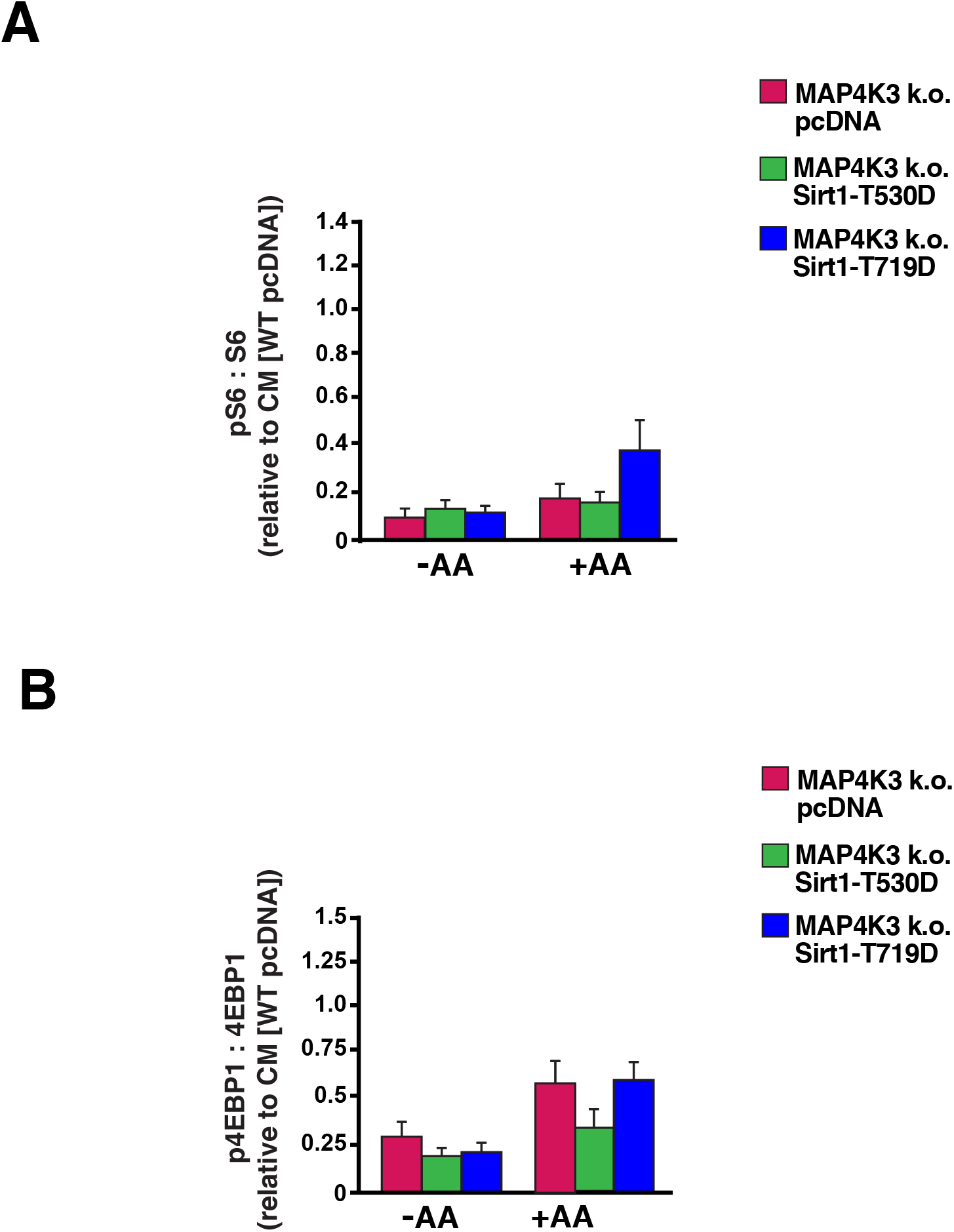
SIRT1-T530D and SIRT1-T719D phosphomimetic mutants are incapable of rescuing mTORC1 activation in MAP4K3 k.o. cells. We transfected MAP4K3 HEK293A k.o. cells with the SIRT1-T530D phosphomimetic mutant, the SIRT1-T719D phosphomimetic mutant, or pcDNA empty vector, and maintained the cells under baseline complete media (CM) conditions, or subjected the cells to amino acid starvation (-AA) or amino acid starvation followed by amino acid restimulation (+AA). As a basis of comparison, we transfected WT HEK293A cells with pcDNA empty vector, and maintained them under baseline complete media (CM) conditions, or subjected the cells to amino acid starvation (-AA) or amino acid starvation followed by amino acid restimulation (+AA). We immunoblotted resulting cell protein lysates for phosphorylated S6, total S6, phosphorylated 4E-BP1, and total 4E-BP1, quantified the levels of phosphorylated S6, total S6, phosphorylated 4E-BP1, and total 4E-BP1 by densitometry, determined the ratio of phosphorylated S6: total S6 and normalized the results to pcDNA-transfected WT HEK293A cells at baseline **(A)** or determined the ratio of phosphorylated 4E-BP1: total 4E-BP1 and normalized the results to pcDNA-transfected WT HEK293A cells at baseline **(B)**. *P* = n.s., ANOVA; n = 3 biological replicates. Error bars = s.e.m.

